# SEAMoD: A fully interpretable neural network for cis-regulatory analysis of differentially expressed genes

**DOI:** 10.1101/2023.11.09.565900

**Authors:** Shounak Bhogale, Chris Seward, Lisa Stubbs, Saurabh Sinha

**Affiliations:** Center for Biophysics and Quantitative Biology, University of Illinois at Urbana-Champaign, Urbana, IL 61801, USA; Pacific Northwest Research Insititute, Seattle WA 98122; Wallace H. Coulter Distinguished Faculty Chair in Biomedical Engineering; Wallace H. Coulter Department of Biomedical Engineering, Georgia Institute of Technology, Atlanta, GA 30332; H. Milton Stewart School of Industrial & Systems Engineering, Georgia Institute of Technology, Atlanta, GA 30332

## Abstract

A common way to investigate gene regulatory mechanisms is to identify differentially expressed genes using transcriptomics, find their candidate enhancers using epigenomics, and search for over-represented transcription factor (TF) motifs in these enhancers using bioinformatics tools. A related follow-up task is to model gene expression as a function of enhancer sequences and rank TF motifs by their contribution to such models, thus prioritizing among regulators.

We present a new computational tool called SEAMoD that performs the above tasks of motif finding and sequence-to-expression modeling simultaneously. It trains a convolutional neural network model to relate enhancer sequences to differential expression in one or more biological conditions. The model uses TF motifs to interpret the sequences, learning these motifs and their relative importance to each biological condition from data. It also utilizes epigenomic information in the form of activity scores of putative enhancers and automatically searches for the most promising enhancer for each gene. Compared to existing neural network models of non-coding sequences, SEAMoD uses far fewer parameters, requires far less training data, and emphasizes biological interpretability.

We used SEAMoD to understand regulatory mechanisms underlying the differentiation of neural stem cell (NSC) derived from mouse forebrain. We profiled gene expression and histone modifications in NSC and three differentiated cell types and used SEAMoD to model differential expression of nearly 12,000 genes with an accuracy of 81%, in the process identifying the Olig2, E2f family TFs, Foxo3, and Tcf4 as key transcriptional regulators of the differentiation process.

## INTRODUCTION

A popular approach to the study of molecular mechanisms underlying any biological process involves transcriptomic profiling and determination of differentially expressed (DE) genes, followed by analysis of their promoters for potential transcriptional regulatory mechanisms (1–3). As cis-regulatory signals are often present not only in promoters but also in more distally located enhancers, it is common to use genome-wide epigenomic profiling, e.g., using ATAC-seq (4) or ChIP-seq (5) assays, to identify candidate enhancers that may then be searched for motifs (6). With a set of DE genes identified and with knowledge of their associated promoters or enhancers, one may use statistical (7, 8) or machine learning (9) tools to discover over-represented or discriminative motifs in the regulatory sequences and compare these motifs with compendia of known motifs such as JASPAR (10) and CIS-BP (11) to implicate TFs in the regulation of those genes. Alternatively, one may employ available tools to directly test for association between known transcription factor (TF) motifs from existing compendia and the regulatory sequences of DE genes (12–14). In either case, one obtains a set of TFs that may underlie the transcriptional changes observed, which may provide various paths forward for detailed mechanistic investigations.

Going beyond identification of TF motifs associated with DE genes, one seeks to quantify the importance of each identified motif to the transcriptional changes overall, or to the regulation of individual DE genes. This may be achieved via models that relate the presence (or strength) of TF motifs in a regulatory sequence to the associated gene’s expression, the so-called “sequence-to-expression” models (15–17). Various approaches have been proposed for such modeling, using techniques from machine learning (18) or adopting biophysical principles (19) to quantify the relative contributions of different TF motifs. Typically, such models require the relevant motifs to be known *a priori* and do not attempt to discover them *ab initio*. Alternatively, one may approach sequence-to-expression modeling without explicitly utilizing TF motifs, e.g., using *k-*mers in the sequence (9, 20, 21) or scanning with convolutional filters (17), but these approaches are primarily concerned with the expression prediction task itself and not the discovery and relative importance of TF motifs.

Here, we present a computational tool for cis-regulatory analysis that (1) discovers TF motifs from candidate regulatory sequences (enhancers), (2) predicts each enhancer’s regulatory potential from the combination of motifs present therein and (3) relates that regulatory potential, along with epigenomic information about the enhancer, to differential expression of the associated gene, all in one simultaneously learned model. This tool, called SEAMoD (**S**equence-, **E**xpression-, and **A**ccessibility-based **Mo**tif **D**iscovery), implements a fully interpretable neural network to relate enhancer sequences to differential gene expression. Additionally, it can make use of epigenomic information provided in the form of candidate enhancers for each gene, with associated scores reflecting local chromatin accessibility, and automatically search for the most promising enhancer among the candidates. Furthermore, SEAMoD is a multi-task learner capable of examining differential expression associated with multiple biological conditions, such as several differentiated cell types compared to a progenitor cell type, thus sharing information across the different conditions in its search for underlying TF motifs. While a number of previous studies have used deep neural networks to map sequences of segments to “functional” readouts such as epigenomic states (22–29), SEAMoD focuses specifically on prediction of differential gene expression between conditions, uses far fewer parameters and thus requires far less training data, and emphasizes biological interpretability.

The inputs to SEAMoD include (bulk) transcriptomic and epigenomic data on two or more biological conditions or cell types, one of which is the reference and the others are called target conditions: the transcriptomic information is in the form of set of DE genes (up- or down-regulated) in each target condition relative to the reference condition, and the epigenomic information is in the form of a set of candidate enhancers for each gene in each condition, and their regulatory activity scores, e.g., from DNA accessibility (30) or H3K27ac ChIP-seq (31) assays. SEAMoD automatically selects one candidate enhancer per condition (for each gene), uses a large set of learnable convolutional filters (representing TF motifs) to parse each enhancer for its motif content, then predicts the enhancer’s condition-specific regulatory potential based on its combinatorial motif content. Next, it scales this regulatory potential by the enhancer’s measured activity in that condition, and predicts differential gene expression by comparing regulatory potentials in the target and reference conditions. Its outputs include the learned TF motifs in position weight matrix (PWM) form and regulatory role (strength, directionality) of each motif in the transcriptomic changes of each target condition relative to the reference.

We used SEAMoD to study the transcriptional regulatory mechanisms underlying neural stem cell (NSC) differentiation *in vitro*. We cultured NSCs from mouse embryonic forebrain, differentiated these into neurons, astrocytes and oligodendrocytes and profiled these differentiated cell types, as well as the undifferentiated NSCs, using RNA-seq and H3K27ac ChIP-seq. Treating the differentiated cell types as target conditions and NSC as the reference condition, we used SEAMoD to identify TF motifs related to expression changes accompanying the differentiation processes, including Olig2, E2f family TFs, Foxo3, Tcf4, and others, for which there are varying levels of literature evidence supporting their role in neuronal differentiation. The SEAMoD predicted roles (activator or repressor) of identified TFs were by and large consistent with the literature, where such information was available.

## METHODS

### Establishment and maintenance of mouse embryonic neural stem cells

Pregnancies were identified in C57BL/6J females by detection of vaginal plugs (day of plug detection is E0.5), and embryos were extracted at E14.5. Whole embryonic brains were transferred to a sterile petri dish containing ice cold PBS with 2% glucose and frontal cortices were separated essentially as described (32). Collected tissues were then transferred to a 15 mL conical tube containing Complete Embryonic NeuroCult Proliferation Medium (NeuroCult Basal Medium (Cat. No. 05700) containing NeuroCult Proliferation Supplement (Cat. No. 05701) with 20 ng/mL rh EGF (Cat. No. 02633) (Stemcell Technologies, Cambridge, MA), 1X Pen Strep (Life Technologies, Carlsbad, CA) and triturated with a pipette to acquire single-cell suspension. Clumps including connective tissues and undissociated cells were excluded by allowing the solution to settle for 3 minutes. Supernatant was then transferred to a 15 mL conical tube containing 10 mL of Complete Embryonic NeuroCult Proliferation Medium (Stemcell Technologies, Cambridge, MA) and centrifuged at 500 × g for 5 minutes. Cell pellets were resuspended with Complete Embryonic NeuroCult Proliferation Medium and counted using hemocytometer. Approximately 2.0 × 10^6^ cells were plated to T-25 flasks which were previously coated with 10 ug/mL poly-D-lysine (PDL; Sigma-Aldrich, Cat. No. P7280) and 10 ug/mL laminin (LMN; Sigma-Aldrich, Cat. No. L2020) and incubated at 37° C with 5% CO2. Complete culture medium was replaced with fresh medium the following day.

### Neuron, oligodendrocyte, astrocyte-specific differentiation

Approximately 5.0 × 10^6^ of proliferative mouse neural stem cells were plated to PDL/LMN-coated T-75 flasks. Mouse neuron differentiation medium (Neurobasal medium with 2% B-27 supplement and 2 mM GlutaMAX-1 supplement [Life Technologies, Carlsbad, CA, otherwise mentioned]), oligodendrocyte differentiation medium (Neurobasal medium with 2% B-27 supplement, 2 mM GlutaMAX-1, and 30 ng/mL T3), and astrocyte differentiation medium (Dulbecco’s Modified Eagle Medium with 1% N-2 supplements, 2 mM GlutaMAX-1, and 1% Fetal Bovine Serum) were freshly prepared before use. Upon reaching 70% confluency, complete culture medium was replaced with corresponding cell-type specific differentiation medium after two 1X PBS washes. Differentiation media was changed every 48 hours and samples were collected at div 0, div 2, and div 4.

### RNA Extraction and Library Preparation

About 0.5 × 10^7^ cultured cells were disrupted in Trizol (Life Technologies, Carlsbad, CA, USA) with a cell scraper. The aqueous phase was precipitated in isopropanol, resuspended in nuclease-free water, treated with DNase I (New England Biolabs, Ipswich, MA, USA), and cleaned up using a Zymo RNA Clean & Concentrator™-25 kit (Zymo Research, Irvine, CA, USA) according to manufacturer’s specifications. Prior to library preparation, total RNA was checked for purity using a NanoDrop ND-1000 spectrophotometer, integrity using RNA Nano chips on an Agilent 2100 Bioanalyzer, and concentration using a Qubit 2.0 fluorometer. RNA-Seq libraries were prepared from total RNA robotically using TruSeq Stranded mRNA HT (Illumina, San Diego, CA, USA) on an epMotion 5075 robot (Eppendorf, Hamburg, Germany). Libraries were pooled and sequenced on an Illumina HiSeq 2500 sequencer by the UIUC W. M. Keck Biotechnology Center using an Illumina TruSeq SBS sequencing kit, version 3 (Illumina, San Diego, CA, USA). All samples were sequenced in single end format with fragment length of 100 bp. Base calling and demultiplexing into FASTQ files was done using CASAVA version 1.8.2 (Illumina, San Diego, CA, USA). Read depth ranged from 45m reads to 65m reads per sample.

### Chromatin immunoprecipitation and library preparation

Chromatin immunoprecipitation was carried out as essentially as described (Saul et al., 2017; Seward et al., 2022). About 2.0 × 10^6^ cells were fixed in PBS with 1% formaldehyde for 10 minutes. Fixing reaction was stopped with addition of glycine to 0.125M. Fixed cells were washed 3x with PBS+Protease inhibitor cocktail (PIC, Roche) to remove formaldehyde. Washed cells were lysed to nuclei with lysis solution – 50 mM Tris-HCl (pH 8.0), 2 mM EDTA, 0.1% v/v NP-40, 10% v/v glycerol, and PIC – for 30 minutes on ice. Cell debris was washed away with PBS with PIC. Nuclei were pelleted and flash-frozen on dry ice. Cross-linked chromatin was prepared and sonicated using a Bioruptor UCD-200 in ice water bath to generate DNA fragments 200-300 bp in size. 2mg of Rabbit polyclonal antibody raised against acetylated histone H3 lysine 27 (Abcam ab4729) was used to precipitate chromatin from nuclei of approximately 1 × 10^6^ cells. DNA was released and quantitated using Qubit 2.0 (Life Technologies) with dsDNA HS Assay kit (Life Technologies, Q32854), and 10 ng of DNA was used to generate libraries for Illumina sequencing. ChIP-seq libraries were generated using KAPA LTP Library Preparation Kits (Kapa Biosystems, KK8232) to yield two independent ChIP replicates for each antibody. We also generated libraries from sonicated genomic input DNA from the same chromatin preparations as controls. Libraries were bar-coded with Bioo Scientific index adapters and sequenced to generate 20-40 million reads per duplicate sample using the Illumina Hi-Seq 2500 instrument at the University of Illinois W.M. Keck Center for Comparative and Functional Genomics using a HiSeq SBS sequencing kit version 4 according to manufacturer’s instructions. Fastq files were generated and demultiplexed with the bcl2fastq v2.17.1.14 Conversion Software (Illumina).

### RNA-Seq Bioinformatics

FASTQ files were aligned to the Ensembl annotation of the NCBIM37 version of the mouse genome using TopHat2 version 2.0.8 (33). Reads inside of exon features were counted in union mode using htseq version 0.6.1 (34). Differential expression analysis was done in R using the Bioconductor package edgeR (35) after filtering for genes with expression ≥ 1 CPM within the smallest group for pairwise comparisons. Pairwise comparisons were analyzed using the bin.loess version of trended dispersion in edgeR version 3.2.4 in R version 3.0.0, essentially as previously described (36).

### ChIP-Seq Bioinformatic analysis

ChIP-seq reads were mapped to the mm9 mouse genome build with bowtie2 and H3K27Ac peaks were identified using the HOMER software suite v4.7 (37), as previously described (36, 38). Peaks were called with default settings, except local filtering was disabled (-L 0), peak size was set at 200bp, and minimum distance between peaks was set at 150bp. 1001bp FASTA sequences were extracted around each peak center using HOMER. Sequencing data has been deposited in the GEO archive under GSE236450. Chromatin profiles are available online as a UCSC Genome Browser track hub at https://trackhub.pnri.org/stubbs/ucsc/public/culture.

### Bioinformatics Data Pre-processing

We focused on a subset of genes that pass certain filters including the fold change and FDR for differential expression and also for having at least one H3K27Ac peak within 100kb of the gene’s transcription start site (TSS), as outlined next.

First, we calculated the fold change in the expression levels for a gene in the reference cell type (NSC) and the target cell types (Neurons, Astrocytes, and Oligodendrocytes). A gene was called differentially expressed in the target cell type if the fold change was at least 20% and the FDR was <= 0.1. All remaining genes were assigned the “no-change” class. Next, we removed all those genes which were not differentially expressed in any of the target cell types, thus ensuring that each gene in the analyzed set was differentially expressed in at least one of the three cell types.

We also limited the genes analyzed based on the presence of potential cis-regulatory elements in their proximity. For this, we first counted the number of H3K27Ac peaks within the 100kb of the TSS for a gene. Note that since chromatin landscape is not uniform, we restricted the 200kb window (100kb in each direction of TSS) to a window of smaller length if it overlapped with a TAD boundary (31) or inaccessible region such as telomeres and centromeres. We removed all the genes that did not have any H3K27Ac peak in the 200kb window in all cell types. Thus, the remaining genes had at least one H3K27Ac peak in each of four cell types. These criteria, involving differential expression and presence of H3K27ac peaks, yielded 11,622 genes in total.

Each gene had a set of candidate enhancers (H3K27ac peaks) specific to each cell type, with some genes surrounded by a large number of peaks around the TSS while others only had a few (Supplementary Figure 1). For uniformity, we considered the nearest four peaks as candidate enhancers for a gene-cell type pair. Thus, a gene could at the most have 16 candidate enhancers (four in each cell type).

### Neural network model and training

The model is designed to predict the differential expression of a gene in each target condition relative to the reference condition, based on sequences of candidate enhancers in all conditions (one per condition) and their condition-specific activity scores (Figure 1B). Each enhancer sequence is represented by 1-hot encoding, thus a 4 × *L* matrix where *L* is the enhancer length. (All enhancers are of same length.)

**Figure 1:**
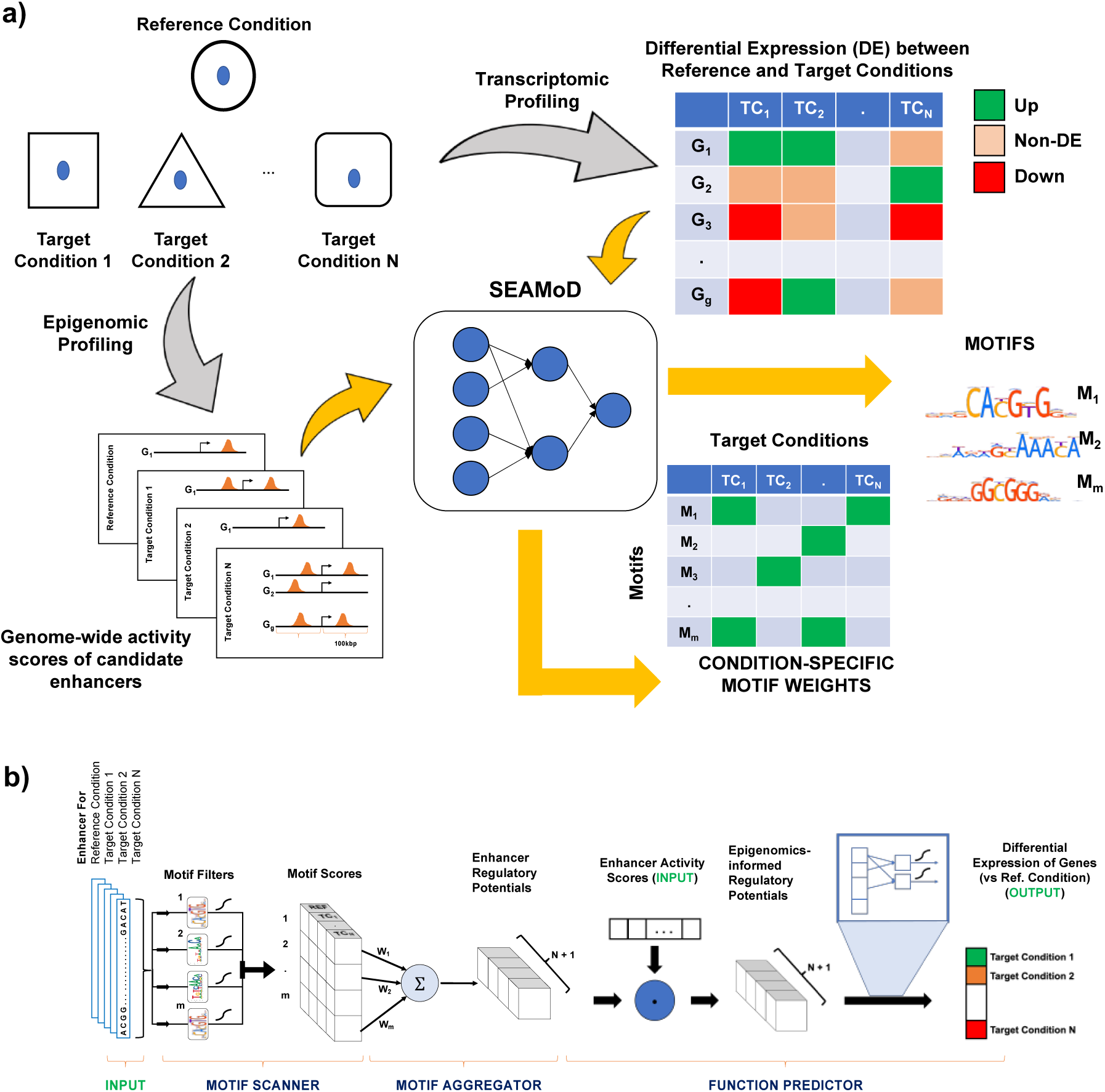
**a) Input/Output of SEAMoD** Inputs include data on one or more “target conditions” and a “reference condition”. Transcriptomic data inputs are in the form of genes that are up- or down-regulated in each target condition vs. reference condition. Epigenomic data input are in the form of candidate enhancers for each gene in each condition, along with their condition-specific levels of activity (e.g., accessibility or H3K27ac mark). From these inputs, SEAMoD trains a neural network model to report TF motifs and their importance (“weights”) to differential expression in each target condition. **b) Model schematic** – The model is comprised of three modules – Motifs Scanner, Motif Aggregator and Function Predictor Module. The first module scans for motif presence in a sequence. The second module calculates a weighted sum of motif scores from the first module, using condition-specific weights, obtaining enhancer regulatory potential scores. The last module scales these regulatory potential scores with activity scores (e.g., accessibility) and uses the resulting scores to predict differential gene expression (up, down or no-change).

#### Motif scanner module

The first module of the neural network comprises a set of convolution filters (called motif filters) with exponential activation, a max pooling layer and an average pooling layer. It processes a condition-specific candidate enhancer to produce a motif presence score for each motif filter. Its function can be written as:

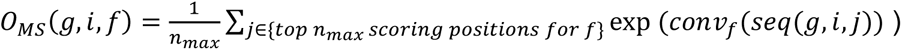

where

- *O*_*MS*_ (*g*, *i*, *f*) = Output of Motif Scanner (MS) module for *i*^*th*^ candidate enhancer of gene *g* for the convolutional filter *conv*_*f*_ (motif presence score),
- *seq*(*g*, *i*, *j*) = *j*^*th*^ position on *i*^*th*^ sequence of gene *g*
- *conv*_*f*_(*seq*(*g*, *i*, *j*)) = output of the convolution of *conv*_*f*_ at *seq*(*g*, *i*, *j*).
- *n*_*max*_ = number of top scoring positions (hyperparameter set by the user)

Informally speaking, the *n*_*max*_ highest scoring matches to the motif represented by a motif filter are averaged to produce the motif presence score. The motif filter *conv*_*f*_ has dimensions 4 *X l* where *l* is the length of the filter (identical across all filters) and is a hyperparameter set by the user. All motif filters are free parameters of the model.

#### Motif aggregator module

This takes the outputs of the motif scanner module (*O*_*MS*_ (*g*, *i*, *f*)) and combines the motif presence scores of a candidate enhancer into a single value, thus generating the “regulatory potential” of the enhancer. Its function may be described as:

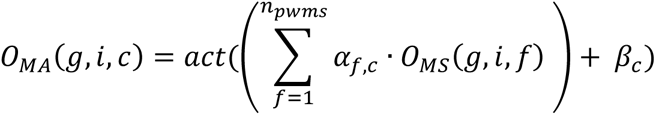

where

- *O*_*MA*_(*g*, *i*, *c*) = output of Motif Aggregator (MA) module for *i*^*th*^ candidate enhancer of gene *g* in the condition *c*, representing the enhancer’s regulatory potential in that condition,
- *act*(∗) = activation function (tanh or ReLU, can be set by user),
- *⍺*_*f*,*c*_ = weight of the motif filter *conv*_*f*_ in the condition *c* (free parameter),
- *m* = number of convolutional (motif) filters (hyperparameter),
- *β*_*c*_ = bias term for condition *c* (free parameter).

#### Function predictor module

This final module processes the regulatory potential scores of enhancers in all conditions (*O*_*MA*_(*g*, *i*, *c*)), along with their condition-specific activity scores, to make the final predictions of differential expression. The output values are the DE status – fold change or direction of change (−1,0, +1) – of the gene in each target condition, relative to the reference condition. Its function may be written as:

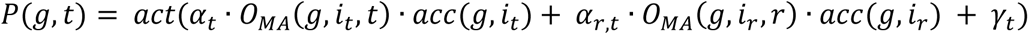

where

- *P*(*g*, *t*) = output (predicted DE) in for gene *g* target condition *t*,
- *act*(∗) = activation function (tanh or ReLU, can be set by user),
- *⍺*_*t*_ = weight for the target condition *t* (free parameter),
- *⍺*_*r*,*t*_ = weight for the reference condition *r* with respect to the target condition *t* (free parameter),
- *i*_*c*_ = index of the candidate enhancer for the condition *c* (*c* ∈ {*r*, *t*}),
- *acc*(*g*, *i*_*c*_) = activity/accessibility score for enhancer with index *i*_*c*_ in the condition *c*,
- *γ*_*t*_ = bias for the target condition *t* (free parameter).

#### Model training

The free parameters of the model include 4*lm* parameters specifying the convolutional (motif) filters in the motif scanner module, (*N* + 1)(*m* + 1) parameters specifying condition-specific motif weights and bias term in the motif aggregator module, and the 3*N* parameters used by the function predictor module. There are three model hyperparameters – *m*, *n*_*max*_, and *l* and three extra hyperparameters involved in training - learning rate and the weights for *L*^1^ and *L*^2^ losses (as weight regularizers). The user may also select the activation function used in the Motif aggregator module and the Function predictor module. We used tanh as the activation function across all layers. The model was trained using ADAM optimizer (39) and weighted mean squared error as the loss function, on the NCSA HAL cluster (40).

The model was first trained using the set of nearest enhancers. This trained model was then used to identify the best enhancer for a gene from the set of four or less enhancers designated for each cell type (mentioned above). Then the model was retrained using the set of these best enhancers for further analysis.

### PWM generation

The motif scanner module was used to identify the best scoring window (with length that of the convolutional filter) from each best enhancer. The best scoring site was discarded if did not pass the ReLU threshold, i.e. if the score of the site was less than zero. A PWM was obtained by taking the average of all the best scoring windows of the corresponding filter. We used ggseqlogo (41) to generate the motif logos from the PWMs.

### Model Code

The entire code is written in python and is available as a github repository which can be accessed by using the following link – https://github.com/shounakbhogale/SEAMoD/

### Motif Matching

We converted the learned convolutional filters into PWMs as mentioned above. Next we used publicly available PWM comparison tool called TOMTOM (42) to compare our PWMs with the known mouse motifs in the HOCOMOCO human and mouse motif database (available in the tool).

### Random baseline accuracy

We used two random baselines where the classifier predicts a gene as up-regulated, non-DE, or down-regulated at random. (Predictions of DE status in different target conditions are made independently.) The uniform random classifier predicts each of the three classes with a probability of 1/3 and thus has an expected accuracy of 1/3. The size-aware random classifier assigns a gene to a class based on the proportion of that class in the data set. Thus, if there are *n*_*i*_ genes of class ‘*i*’ and the total number of genes is *N*, then, this classifier will make the call ‘*i*’ with probability *p*_*i*_ = *n*_*i*_/*N*, with an expected accuracy of 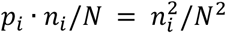 of those calls, and an overall expected accuracy of 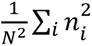. This expected accuracy depends on the class sizes for each target condition, and the average over all three target conditions was 0.365.

## RESULTS

### An interpretable neural network for discovery of motifs underlying differential expression

We developed a computational tool called SEAMoD (**S**equence-, **E**xpression-, **A**ccessibility- based **Mo**tif **D**iscovery) that models differential expression of genes in terms of motifs present in regulatory regions of those genes. It utilizes user-provided genome-wide data on accessible or active chromatin regions to narrow its search for relevant cis-regulatory elements, and it discovers the relevant motifs automatically during model fitting.

The intended application scenario of SEAMoD is the following (**Figure 1A**). Assume we have expression and epigenomic data for one experimental condition or cell type (henceforth called the “reference condition”) and one or more (say *N*) additional conditions or cell types (henceforth called the “target conditions 1 … *N*”). The expression data may be of any type (e.g., RNA-seq) that allows determination of differentially expressed (DE) – up-regulated or down-regulated – genes between each target condition and the reference condition. The epigenomic data may also be of any type, as long as they provide a set of candidate enhancers (e.g., DNA accessibility or H3K27ac ChIP-seq peaks) genome-wide, for each condition. Given these data, including enhancer sequences, SEAMoD discovers a set of position weight matrix (PWM) motifs and uses the presence of these motifs in candidate enhancers to predict each gene’s differential expression in each target condition, while also learning the importance of each motif to that condition.

At the heart of SEAMoD is a simple, fully interpretable neural network model (**Figure 1B**) that relates a gene’s candidate enhancers to its DE status. The “inputs” to this model are the sequences of candidate enhancers of the gene, exactly one enhancer per condition (thus *N* + 1 sequences). (Note: we show later how multiple candidate enhancers per gene are handled by SEAMoD.) The input also includes the “activity score” of each enhancer (e.g., DNA accessibility or strength of H3K27ac mark) in all *N* + 1 conditions. The “output” of the neural network is *N* + 1 real numbers, each between −1 and +1 and representing the predicted differential expression of the gene between a target condition and the reference condition. (Values close to −1 or +1 indicate predicted down- or up-regulation respectively, while those close to 0 indicate a prediction of no change in expression.)

The architecture of the neural network comprises three modules, outlined next. The first module, called the “motif scanner”, uses a fixed set of convolutional “filters” (each filter representing a TF motif) to scan the candidate enhancer sequences and score them for motif presence. A predetermined number of such filters (henceforth called “motif filters”), say *m*, is allocated, and since the module scores each of the *N* + 1 candidate enhancer sequences with each filter, its output is a *m* × (*N* + 1) matrix of real numbers summarizing the enhancers’ motif profiles. Importantly, the motif filters are free parameters of the model and are learnable from data in the course of model training. The second module of the neural network, called the “motif aggregator”, combines the motif presence scores of an enhancer into a weighted sum that may be interpreted as the regulatory potential of the enhancer, loosely defined as the strength of gene activation (positive values) or repression (negative values) driven by this enhancer in an experimental condition. Weights associated with the *m* motif filters are condition-specific (thus *m* × (*N* + 1) in count) and are learnt from data. Each motif’s weight in a condition may be interpreted as a composite of the cognate TF’s concentration in the condition and its activation/repression potential. Note that this manner of aggregating motif scores to model regulatory potential of an enhancer has been studied in the literature, e.g., Kazemian *et al.*, 2010. The motif aggregator thus computes one real number per condition, i.e., an *N* + 1 dimensional vector of condition-specific regulatory potentials. The third and final module, called the “function predictor”, performs two steps: first, it multiplies the enhancer’s regulatory potential in a condition with its activity score (e.g., accessibility or active chromatin mark) in that condition to obtain an epigenomics-informed regulatory potential, and it then combines these potentials for a target condition and the reference condition to make its final prediction of differential expression of the gene in the target condition. Thus, this module’s (and the model’s final) output is an *N* dimensional vector of DE predictions, one for each target condition.

The learnable parameters of the above model include the motif filters, motif aggregation weights for each condition, and parameters used by the function predictor module. The model learns to predict a gene’s DE status in all target conditions simultaneously, and its use of a common set of motif filters allows motif discovery to exploit the entire data set and not just a pair of conditions at a time. Once the model is trained, the motif filters are converted to the popular PWM format by locating the best scoring match to the filter in each enhancer and combining these matches as if they were binding sites of the cognate TF. SEAMoD also reports the importance of each motif to differential expression in each target condition; we explain this functionality in a later section.

### Transcriptomic and epigenomic profiling of neuronal stem cell differentiation

We sought to use SEAMoD to characterize the transcriptional regulatory mechanisms underlying neuronal stem cell (NSC) differentiation to neurons, astrocytes and oligodendrocytes in the mouse brain. We took advantage of well-established protocols to generate neural stem cell (NSC) cultures from the forebrain of embryonic day 14.5 (E14.5) mouse embryos (44), and to culture the NSC under conditions that favor the differentiation into Neurons, Astrocytes, or Oligodendrocytes, respectively (32). We collected cells at days *in vitro* 0 (undifferentiated NSC), and div4 (initiation of the maturation stage for Neurons, Astrocytes or Oligodendrocytes) to focus on the early stages of cell commitment and differentiation, which are well documented to require the precisely orchestrated activities of conserved transcription factor networks both *in vivo* and *in vitro* (45, 46). We collected RNA and chromatin samples from both stages to generate transcriptomic and histone modification profiles in each of the three types of differentiating cells.

We compared the transcriptomic profiles of the NSC to the three differentiated cell types at div4 (**Figure 2A**), finding a far greater mutual similarity among the differentiated cell types (Pearson correlation 0.84 on average) than between any of them and the stem cells (average correlation 0.61). This is not surprising, given that the cells are not fully differentiated at div 4. It did suggest, however, that many of the underlying regulatory mechanisms (e.g., enhancers and TFs) may be common to the three differentiation paths, since the end points examined are relatively similar to each other. We identified DE genes (FDR 0.1, see Methods) between NSC and each of the differentiated cell types. This yielded a set of 11,622 genes (after filtering them for the epigenomic profiling explained in the next paragraph) that are DE in at least one of the three contrasts, with the frequency of up- and down-regulation in any particular cell type being 39-41% and 40% respectively (**Figure 2B**). (The majority of genes, nearly 63%, are not DE in any cell type and were not included in this figure or in the analyses below.) Among the DE genes, roughly 60% showed the same direction of regulation (up or down) between NSC and each of the three cell types (Supplementary Table 1), consistent with the presence of common regulatory mechanisms across differentiation paths. At the same time, there were 4930 genes with DE status that varies across cell types, suggesting the presence of regulators exclusive to specific differentiation paths.

**Figure 2:**
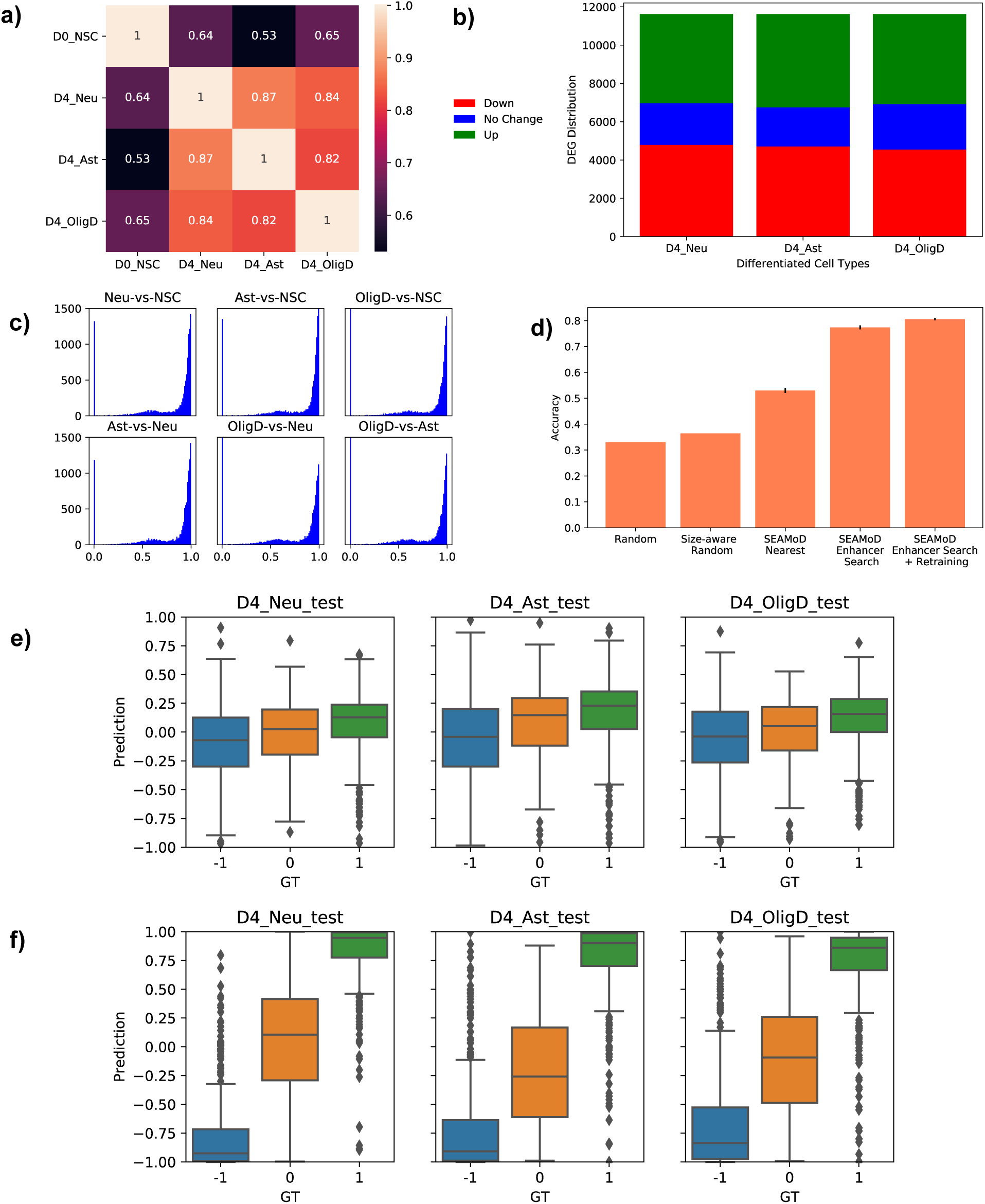
**a)** Gene expression correlation between NSC (D0_NSC) and each of the three differentiated cell types at div4 (neurons – “D4_Neu”, astrocytes – “D4_Ast”, oligodendrocytes – “D4_OligD”). Correlation between differentiated cell types is more pronounced than it is between a differentiated cell type and NSC. **b)** Counts of up- and down-regulated genes as well as non- DE genes in the set of all 11,622 genes that are DE in at least one differentiated cell type relative to NSC. **c)** Histograms of overlaps between the nearest candidate enhancers (H3K27ac ChIP peak) of a gene from pairs of cell types (neurons – “Neu”, astrocytes – “Ast”, oligodendrocytes – “OligD”, stem cell – “NSC”). **d)** Model performance – × axis represents the models (including the two random baselines – “Random” and “Size-aware” random) and Y axis represents the classification accuracy on test set (1,162 genes), averaged over ten folds, along with error bars (standard errors). **e,f)** Raw predicted DE scores of genes in test set, across different target conditions (neurons – “D4_Neu_test”, astrocytes – “D4_Ast_test”, oligodendrocytes – “D4_Olig_test”). Panel (e) shows predictions of SEAMoD in the “nearest enhancer” mode and panel (f) shows the “enhancer search” mode

We restricted the regulatory region of a gene to 100kb on either side of its transcription start site (TSS). We removed genes that do not have any H3K27Ac peak in its regulatory region in any of the four cell types. The average number of peaks in a gene’s regulatory region varied across the four cell types (Supplementary Figure 1) but the nearest peaks showed significant overlaps amongst pairs of cell types (**Figure 2C**, further details below).

### SEAMoD relates sequence to differential expression with high accuracy

Identification of thousands of DE genes related to NSC differentiation led us to investigate their cis-regulatory mechanisms. A common strategy for such investigations is to search for transcription factor (TF) binding sites in active enhancers linked to DE genes (1–3). However, there were two main hurdles in pursuing this strategy. Firstly, we lacked comprehensive prior knowledge of relevant TF motifs or access to relevant TF-DNA binding profiles. Secondly, enhancer-gene linkages, often obtained through “3D chromatin” data (47), were not available for the cellular conditions of our study. To address the latter issue, we initially selected the H3K27ac peak nearest to the transcription start site (TSS) of a gene, in a cell type, as being that gene’s putative enhancer (driver of expression) in the cell type. Each gene was thus assigned four candidate enhancers, one for each cell type. (Each enhancer was delineated as the 1 Kbp segment centered at the summit of the H3K27ac peak.) We observed a very high degree of overlap among these four candidate enhancers for any gene, on average (**Figure 2C**). For instance, the candidate enhancers of any two cell types (e.g., NSC and a differentiated cell type) had a mutual overlap of over 80% for roughly 59-68% of genes. There was also a substantial number of genes (11-17%) for which candidate enhancers from two cell types were non-overlapping (overlap < 20% length). If the candidate enhancers underlie a gene’s differential expression, this meant that DE arises in part from cell type-specific enhancer switching and perhaps in greater part from the same enhancer providing different readouts in different trans environments with varying TF concentration profiles. To investigate cis-regulatory mechanisms more deeply we thus needed a model that accommodates both scenarios. We note that the above inferences are contingent on the “nearest H3K27 peak” strategy yielding the relevant enhancers accurately. This is known to be a sub-optimal strategy, and we will revisit it later.

Having made a preliminary assignment of cell type-specific candidate enhancers for each gene, we next tackled the other major hurdle for cis-regulatory analysis: the discovery of relevant TF motifs. We used the newly developed SEAMoD tool for this purpose, designating the “NSC” as the reference condition and the three differentiated cell types (“Neuron”, “Astrocytes”, “Oligodendrocytes”) as the target conditions, providing it the DE status of each gene in each target condition (versus reference condition), the above-selected candidate enhancers and their condition-specific H3K27ac peak “heights” as activity scores. Recall that SEAMoD simultaneously learns a set of TF motifs and uses their presence in enhancer sequences to predict each gene’s DE status in each target condition. In addition to providing us with relevant motifs and their condition-specific regulatory roles, this has the secondary benefit of allowing us to evaluate the model (and thus the reliability of discovered motifs/roles) through cross validation on genes. We thus divided the full complement of 11,622 genes into “training”, “validation” and “test” sets (in 8:1:1 proportions), using the training and validation sets to fit model parameters and hyperparameters respectively. We evaluated the accuracy of the trained model on the 1,162 genes in the test set, using their known DE status in the three target conditions. Since each predicted status can be −1, 0, or +1 (down-regulated, no-change or up-regulated), a random model is expected to have ∼0.33 accuracy, while a “size-aware” random model, which predicts class labels in proportion to their relative frequencies in the data, has an expected accuracy of ∼0.36. The SEAMoD model showed an accuracy of ∼0.52 on test data (“SEAMoD Nearest” in **Figure 2D**) and ∼0.57 on training + validation data (Supplementary Figure 2), considerably better than the random baselines. The small difference between test and training accuracy also suggested that the model was trained without significant overfitting.

### SEAMoD selects condition-specific enhancers to boost accuracy

A potential limitation of the above approach lies in its use of the “nearest H3K27ac peak” strategy to associate candidate enhancers with genes. To address this, SEAMoD has an “enhancer search” feature whereby it can examine more than one candidate enhancer per condition for each gene. It first trains the neural network using the candidate enhancers nearest to the TSS, then uses the trained model to predict DE status using every possible combination of candidate enhancers, selecting the combination that yields greatest test accuracy. (The selected combination still includes exactly one candidate enhancer per condition, for a gene, but that enhancer may not be the one nearest to TSS.) Finally, it retrains the previously trained neural network on the selected enhancers.

To use the “enhancer search” feature of SEAMoD we now examined all H3K27ac peaks within 100 Kbp (upstream or downstream) of the TSS of a gene and provided the four nearest peaks as candidate enhancers (for each condition) to the enhancer-search module of SEAMoD. Note that this created a search space of 256 enhancer combinations (for each gene) that were examined by the tool to find the best enhancer combination and retrain the neural network. The accuracy of this final model (“SEAMoD Enhancer Search + Retraining” in Figure 2D) was ∼0.81, a dramatic improvement over the ∼0.52 seen with the “nearest enhancer” strategy. Strictly speaking, this is not a fair comparison since the “enhancer search” results required the test data for enhancer selection, but it nevertheless points to a substantially more accurate sequence-function relationship being captured by the final model.

Recall that the SEAMoD neural network produces a real number between −1 and +1 as its prediction for the gene’s DE status in each condition; this number is then compared with suitable thresholds to output a three-way call of DE status. We examined the real valued predictions made by the neural network when using the nearest enhancer strategy versus the enhancer search strategy (**Figure 2E,F**) and noted that in the latter case (enhancer search mode) the predictions of up- versus down-regulated versus non-DE genes are significantly better separated. Independently, we examined the overlap of a limited reference set of validated enhancers in neural cells (48) with the enhancers used by either strategy and found the enhancer search strategy to exhibit a greater overlap (see Supplementary Table 2). Taken together, these findings support the functional importance of distal enhancers for neuronal differentiation, consistent with similar findings in other systems (49). They also provide assurance regarding the quality of the trained models, prompting us to look closer into their functioning.

### From model to motifs

The final model trained with the enhancer-search strategy included 128 motif filters, which are reported by SEAMoD in PWM format (Supplementary File S1). For mechanistic interpretation, the tool ranks these motifs by their relevance to each target condition and to all conditions in aggregate. This is achieved by nullifying the contribution from a motif filter to the model’s predictions for all genes for a target condition (or all target conditions) and assessing the resulting decrease in model accuracy when predicting DE status in that condition (or all conditions); see Methods. The assessed deterioration in performance then serves as a numeric score of the motif’s relevance to each condition or for overall relevance. Motifs are reported ranked by this score.

We focused our attention to the top 10% (13 of 128) motifs for each target condition and used the resulting set of 27 motifs (**Figure 3A**) for further examination. Several of these motifs contributed 3% or more to the overall accuracy of the model (column ‘Total’). A few of these 27 motifs were relevant to all three differentiated cell types while several were important only to individual cell types (**Figure 3A**), pointing to a mix of shared and condition-specific regulatory mechanisms.

**Figure 3:**
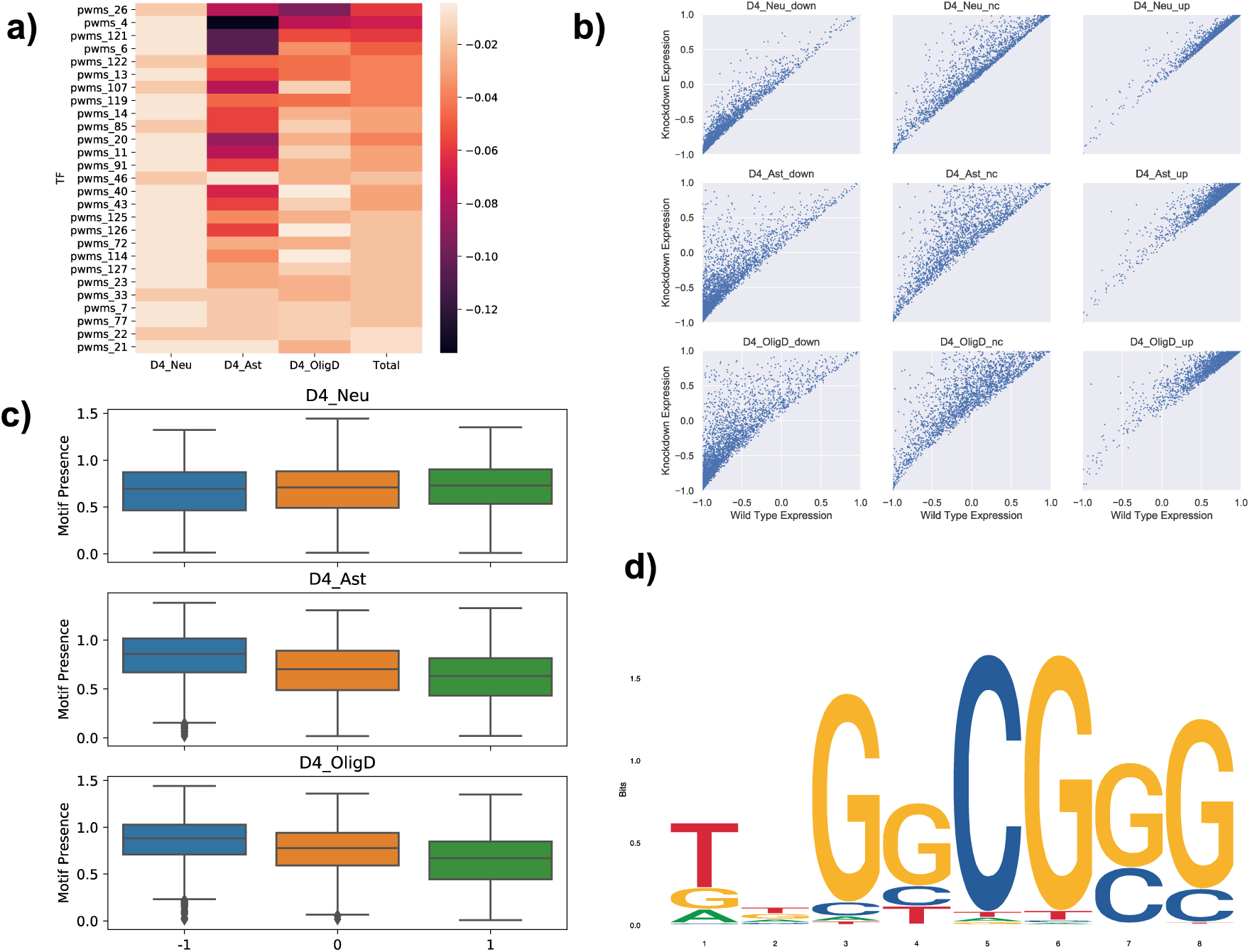
**a)** TF motif importance for neuronal differentiation: Heatmap represents the change in accuracy of the model when a TF motif’s contribution is made zero while keeping the rest of the parameter values fixed. Each row represents a convolution filter (TF motif) whereas the columns show importance to each of the three differentiated cell types and to all cell types (‘total’). The topmost row (“pwm_26”) has the highest overall importance (‘total’) among the 128 pwms learnt by the model. **b)** DE scores of all genes, predicted by the wildtype model (X axis) and the model with pwm_26 “knockdown” (Y axis). Each subplot represents a differentiated cell-type and DEG class. The filter acts as a repressor in the model as all points lie above or on the diagonal. **c)** Motif presence scores for the pwm_26 motif in all enhancers selected by the SEAMoD enhancer search mode. The three panels show scores for enhancers selected for each of the target conditions (Neurons – ‘Neu’, Astrocytes – ‘Ast’, Oligodendrocytes – ‘OligD’) and categorized by the direction of regulation of the associated gene (up – ‘1’, no-change – ‘0’, down – ‘-1’). **d)** pwm_26 logo: The motif is obtained by averaging the best scoring matches to the convolution filter in each enhancer.

The motif with the greatest overall importance is “pwms_26” (**Figure 3D**), which matches known E2F family motifs. Nullifying its contribution reduces the overall accuracy of the model from 84% to 78% (on the entire dataset), the largest such effect among all motifs, and its importance score is ranked 1^st^, 7^th^ and 1^st^ for Neurons, Astrocytes and Oligodendrocytes respectively. **Figure 3B** shows the predicted differential expression of each gene with the full model (“wild type”, × axes) and upon nullifying this motif (“knockdown”, y axes), grouped by the true DE status of genes. Removing this motif from the trained model results in greater predicted DE scores, i.e., the model uses the motif as a repressive influence on gene expression in each differentiated cell type (or equivalently, as an activating influence in the NSC). In fact, for 5,272 gene-condition combinations the predicted DE status changes (from down-regulated to non-DE or non-DE to up-regulated) upon removal of this motif. The motif’s effect is more pronounced for the Oligodendrocyte and Astrocyte down-regulated genes. A direct examination of the strength of this motif’s presence in the optimal enhancers linked to different DE gene classes (**Figure 3C**) also revealed a significant enrichment in down-regulated genes (compared to up-regulated genes) of these two cell types. We discuss biological significance of this and other motifs in the next section.

### A shortlist of putative transcription factors underlying NSC differentiation

We obtained above a list of 27 motifs that are relevant to at least one of the differentiated cell types. These motifs were by and large specific (moderate to high information content) and non-redundant. We compared these motifs to experimentally characterized TF motifs in mouse, using the TOMTOM software (42), and noted matching TFs (**Figure 4**) that were also supported by high expression in the relevant cell types according to our expression data (see Methods). (For 11 of these motifs no TF matches meeting these criteria were found.)

**Figure 4:**
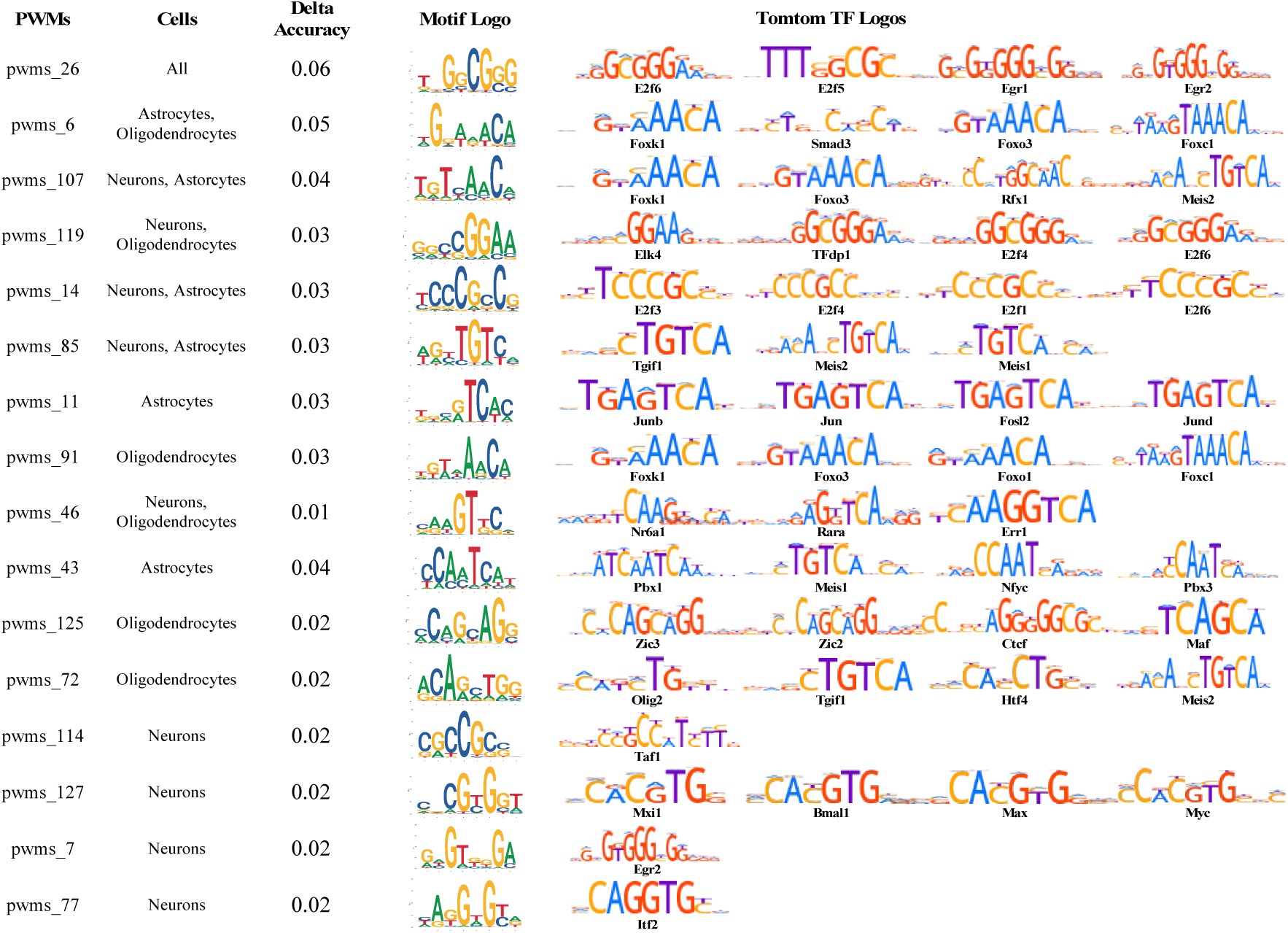
Important motifs identified by the model. The model was trained with 128 convolutional filters. The filters were then converted to PWMs. Only 16 of them survive various criteria including the effect of in-silico knockdown on the overall accuracy, TOMTOM motif matches and their strengths, and expression thresholds (Methods). The first column in the table represents the motif identifier, second column represents the cell types that the motif was found to be important to, third column represents the change in total accuracy upon nullifying the motif’s contribution, and the last column lists the significant and highly expressed TOMTOM TF motif matches. Note: in cases where a PWM matches more than four known TF motifs, only four matches have been shown.

The TF Olig2 has is a match to pwm_72, which was found important to DE status prediction in Oligodendrocytes. It is one of the well-known TFs involved in and also required for Oligodendrocyte differentiation. In fact, it is at the top of the TF hierarchy that controls Oligodendrocyte development (50–52). Another set of TFs that had matching pwms learnt by the model are E2f family of proteins (pwm_14, pwm_26 and pwm_119, Figure 4). E2f proteins are required for proliferation of neuronal precursors and inhibit cell differentiation (53), and our cells were exactly in this phase of the differentiation. Furthermore, all three corresponding motifs were learnt as repressors and there is strong evidence in the literature that this family of proteins mostly act as repressors by interacting with other co-repressors. Another TF that had multiple matching pwms is Foxo3 (pwms_6, pwms_91, and pwms_107, **Fig 4**). It is a well-known inhibitor of neuron differentiation and it also maintains undifferentiated progenitor pools and so it is expected to be important in all the three differentiated cell types (54–56). TCF4 (also known as Itf2), a central TF involved in neuronal differentiation (57) had a matching PWM (pwms_77) which was indeed found to be important to differential expression in Neurons in the knockdown analysis. Another important neuronal TF that had a matching PWM is Mxi1 (pwms_127). It is essential for neurogenesis and is also an important Myc co-factor (58). The same PWM also matched with other neuronal TFs including Bmal1, Myc, and Max all of which have literature evidence for being involved in neurogenesis (59–61). Note that even though this PWM matched with motifs of multiple TFs, most of these TFs have literature evidence of being important in the relevant cell type (which is neurons in the case of pwms_127). The remaining TFs that had matching PWMs also have some literature evidence that suggest they are important in formation and/or maintenance of these cell types including Zic family proteins, Nr6a1 (62) and Fosl2.

## DISCUSSION

Regulation of gene expression is a complex phenomenon involving multiple transcription factors working in tandem, binding to specific sites in promoters and enhancers that may be located proximal to or distal (tens or hundreds of Kbp) from the regulated gene. The model we presented here uses these regulatory sequences, revealed by epigenomic profiling, to learn the underlying mechanism of differential gene expression between biological conditions or cell types. In the process it learns the relevant sequence signatures (motifs) representing the TF binding sites and also characterizes the effect of the corresponding TFs on gene expression.

One of the salient features of the SEAMoD model is its interpretable architecture. The motif scanner module captures the DNA-binding strengths of TFs, using convolutional filters representing the relevant TF motifs. Note that this module can learn these motifs *ab initio*, but can also use already known PWMs, or fine tune them to fit the data better. The motif aggregator module deals with the effect of a combination of TF binding sites (of varying strengths) on gene regulation, using a simple weighted sum to capture this combined effect. A TF’s regulatory influence depends on two factors (in addition to binding site strengths) – its concentration, which affects its occupancy, and regulatory “potency”, i.e., activation or repression strength of a bound TF (see Dibaeinia and Sinha, 2021). The weight learnt for each TF motif by the aggregator module combines these two factors into a single weight. Note that each motif has unique weight for a biological condition or cell type, reflecting the condition-dependent concentrations and possibly context-dependent regulatory potency of the TF. This interpretability allows us to make meaningful observations about a gene regulatory program and its mechanisms.

Motif discovery is a well-studied problem, and several approaches have been proposed for identifying motifs overrepresented in a set of sequences (64) or from TF-DNA binding data (65). *Ab initio* motif finding methods such as these are generally not designed for explicitly modeling gene expression (or differential gene expression), although there has been at least one effort (MatrixREDUCE (66)) at discovering motifs from a combination of gene expression data and promoter sequences. SEAMoD offers a greater versatility of application scenarios than MatrixREDUCE, allows use of epigenomics profiles, and handles multiple motifs simultaneously.

On the other hand, modeling of gene expression from promoter or enhancer sequences is also an intensely researched problem, but such modeling efforts use pre-specified TF motifs and do not attempt to learn them as part of the model. Xpresso (17) is a state-of-the-art method for modeling gene expression from sequences that does not require pre-specified motifs, but it only allows analysis of promoter sequences.

Neural network models have gained wide popularity for sequence modeling (70). These models, e.g., DeepSEA (23), Enformer (28), DanQ (25), Bassenji (26), relate the sequence of a genomic segment to various functional measurements at that segment, such as chromatin states, TF binding, etc. They are usually set up as multi-task models that learn to simultaneously predict multiple (often hundreds of) types of measurements at a genomic segment and rely on such extensive genomic profiling (many “tracks”) to train their large numbers of parameters. SEAMoD is far smaller and simpler model of sequence interpretation, designed for a much more specific goal: to predict differential expression of a gene associated with an enhancer. It only requires DE status information (at the gene level) and candidate enhancers, which could be derived from a single type of epigenomic profile (H3K27ac in our case), yet achieves strong predictive accuracy for its task. In fact, our primary design criterion in developing SEAMoD was that of full biological interpretability, and predictive accuracy was a supporting goal. Admittedly, SEAMoD in its current version lacks the ability to incorporate multiple genome-wide “tracks” of data that larger models use to their advantage. As such, it may not be ideally suited for certain biological systems and conditions, e.g., the ENCODE-selected cell lines (31), where extensive profiling has already been performed, but this apparent weakness becomes a strength for systems for which such data are lacking. We note that the simple architecture of SEAMoD will facilitate future work on addition of modules that can use additional regulatory datasets mentioned above. Another line of future research related to SEAMoD is to improve the enhancer search module, which is currently a brute-force search, and may in the future be replaced with sampling or greedy strategies (71). In conclusion, we believe that the neural network architecture of SEAMoD provides a good starting point for the family of models that are fully interpretable yet allow for the flexibility that is associated with the deep-learning models.

## DATA AVAILABILITY

Sequencing data has been deposited in the GEO archive under GSE236450. Chromatin profiles are available online as a UCSC Genome Browser track hub at the following link - https://trackhub.pnri.org/stubbs/ucsc/public/culture.

The code is available as a github repository at - https://github.com/shounakbhogale/SEAMoD

## ACKNOWLEDGEMENTS AND FUNDING

We thank Prakruthi Burra for her assistance in setting up the initial framework of the code, Payam Dibaeinia for his expert advice on deep learning models. We also thank Younguk Sun for establishing the NSC culture methods, Huimin Zhang for expert advice and assistance with ChIP, and Joseph Troy for advice and assistance with RNA-seq informatics. This work was supported by a grant from the National Institutes of Mental Health, R01 MH114600 (awarded to LS) and a grant from National Institutes of Health, R35GM131819A (awarded to SS).

