## Supplementary S1 for "SEAMoD: A fully interpretable neural network for cis-regulatory analysis of differentially expressed genes"

```
>pwms_0 8
0.001	0.916	0.003	0.08
0.135	0.854	0.0	0.011
0.093	0.571	0.009	0.328
0.024	0.433	0.516	0.027
0.004	0.798	0.134	0.064
0.475	0.508	0.002	0.015
0.0	0.999	0.0	0.001
0.001	0.997	0.001	0.001
<
>pwms_1 8
0.008	0.954	0.038	0.001
0.008	0.013	0.963	0.016
0.281	0.021	0.0	0.698
0.127	0.022	0.75	0.101
0.161	0.169	0.498	0.171
0.523	0.232	0.158	0.087
0.002	0.569	0.369	0.06
0.567	0.008	0.001	0.424
<
>pwms_2 8
0.802	0.041	0.128	0.029
0.967	0.0	0.033	0.0
0.396	0.0	0.38	0.223
0.043	0.691	0.264	0.002
0.491	0.0	0.007	0.502
0.704	0.042	0.253	0.001
0.531	0.146	0.302	0.022
0.019	0.895	0.083	0.003
<
>pwms_3 8
0.11	0.039	0.848	0.003
0.0	0.003	0.995	0.002
0.03	0.0	0.884	0.085
0.0	0.012	0.948	0.041
0.29	0.013	0.55	0.146
0.133	0.54	0.168	0.158
0.102	0.02	0.009	0.869
0.313	0.152	0.469	0.065
<
>pwms_4 8
0.935	0.003	0.0	0.062
0.328	0.161	0.03	0.481
0.042	0.002	0.57	0.386
0.183	0.159	0.385	0.272
0.076	0.14	0.681	0.103
0.315	0.045	0.003	0.637
0.001	0.412	0.028	0.559
0.964	0.0	0.024	0.012
<
```

```
>pwms_5 8
0.641	0.334	0.02	0.005
0.081	0.048	0.075	0.796
0.605	0.039	0.078	0.278
0.037	0.263	0.022	0.678
0.032	0.395	0.514	0.059
0.05	0.907	0.039	0.003
0.511	0.002	0.29	0.197
0.027	0.836	0.006	0.13
<
>pwms_6 8
0.4	0.106	0.038	0.455
0.003	0.0	0.997	0.0
0.132	0.075	0.557	0.236
0.65	0.002	0.0	0.348
0.463	0.036	0.396	0.105
0.743	0.0	0.018	0.238
0.0	0.913	0.061	0.026
0.848	0.09	0.038	0.024
<
>pwms_7 8
0.263	0.005	0.686	0.046
0.365	0.31	0.0	0.325
0.006	0.001	0.992	0.001
0.144	0.055	0.006	0.795
0.0	0.187	0.432	0.38
0.096	0.107	0.579	0.218
0.032	0.0	0.967	0.002
0.803	0.184	0.011	0.001
<
>pwms_8 8
0.0	0.825	0.165	0.01
0.966	0.012	0.0	0.022
0.017	0.089	0.788	0.106
0.0	0.941	0.059	0.0
0.372	0.295	0.041	0.293
0.019	0.785	0.197	0.0
0.516	0.005	0.343	0.137
0.0	0.0	0.982	0.018
<
>pwms_9 8
0.262	0.333	0.075	0.329
0.011	0.978	0.004	0.007
0.0	0.001	0.999	0.0
0.01	0.614	0.031	0.346
0.026	0.035	0.812	0.126
0.011	0.73	0.003	0.255
0.202	0.026	0.615	0.157
0.211	0.261	0.192	0.335
<
```

```
>pwms_10        8
0.017   0.115   0.0     0.867
0.1     0.435   0.124   0.342
0.812   0.001   0.079   0.108
0.902   0.034   0.065   0.0
0.516   0.011   0.276   0.197
0.783   0.005   0.199   0.012
0.319   0.119   0.01    0.552
0.064   0.004   0.847   0.085
<
>pwms_11        8
0.116   0.016   0.401   0.467
0.143   0.075   0.465   0.318
0.416   0.434   0.086   0.064
0.401   0.016   0.583   0.0
0.021   0.004   0.008   0.968
0.0     0.969   0.001   0.03
0.68    0.002   0.004   0.314
0.231   0.721   0.001   0.047
<
>pwms_12        8
0.0     0.925   0.073   0.002
0.414   0.002   0.17    0.415
0.084   0.0     0.907   0.009
0.182   0.817   0.0     0.0
0.0     0.0     0.0     1.0
0.175   0.199   0.614   0.012
0.0     0.32    0.44    0.239
0.002   0.005   0.802   0.192
<
>pwms_13        8
0.487   0.009   0.16    0.343
0.022   0.669   0.299   0.01
0.0     0.0     0.007   0.993
0.002   0.0     0.841   0.157
0.064   0.0     0.171   0.765
0.28    0.002   0.334   0.384
0.149   0.398   0.182   0.271
0.961   0.012   0.0     0.027
<
>pwms_14        8
0.011   0.275   0.089   0.625
0.042   0.838   0.082   0.038
0.063   0.805   0.103   0.029
0.002   0.951   0.047   0.0
0.121   0.001   0.823   0.055
0.042   0.668   0.185   0.105
0.001   0.965   0.028   0.006
0.069   0.125   0.696   0.11
<
```

```
>pwms_15        8
0.074   0.78    0.084   0.062
0.674   0.256   0.0     0.069
0.091   0.343   0.566   0.0
0.013   0.969   0.001   0.016
0.069   0.035   0.052   0.844
0.041   0.051   0.827   0.081
0.045   0.012   0.937   0.006
0.391   0.59    0.019   0.0
<
>pwms_16        8
0.318   0.224   0.369   0.088
0.583   0.059   0.043   0.315
0.003   0.0     0.992   0.005
0.122   0.05    0.231   0.597
0.016   0.0     0.981   0.003
0.796   0.0     0.036   0.167
0.899   0.0     0.101   0.0
0.437   0.001   0.198   0.364
<
>pwms_17        8
0.079   0.123   0.032   0.766
0.003   0.005   0.008   0.984
0.244   0.0     0.75    0.006
0.035   0.032   0.0     0.933
0.002   0.369   0.105   0.525
0.312   0.187   0.024   0.476
0.463   0.078   0.007   0.452
0.014   0.093   0.537   0.355
<
>pwms_18        8
0.4     0.001   0.279   0.319
0.0     0.001   0.001   0.998
0.005   0.027   0.968   0.0
0.272   0.024   0.703   0.0
0.345   0.197   0.139   0.319
0.181   0.03    0.0     0.789
0.0     0.574   0.075   0.351
0.0     0.282   0.078   0.64
<
>pwms_19        8
0.101   0.837   0.0     0.062
0.14    0.031   0.008   0.821
0.008   0.083   0.908   0.0
0.227   0.023   0.455   0.294
0.058   0.718   0.186   0.038
0.092   0.905   0.003   0.0
0.004   0.0     0.0     0.996
0.088   0.061   0.625   0.226
<
```

```
>pwms_20        8
0.027   0.169   0.799   0.005
0.586   0.0     0.354   0.06
0.006   0.007   0.905   0.081
0.0     0.0     0.997   0.003
0.056   0.482   0.392   0.07
0.05    0.0     0.0     0.95
0.14    0.445   0.397   0.018
0.694   0.001   0.099   0.206
<
>pwms_21        8
0.345   0.526   0.069   0.06
0.332   0.013   0.462   0.193
0.375   0.029   0.571   0.025
0.022   0.0     0.32    0.658
0.151   0.017   0.83    0.002
0.891   0.004   0.105   0.0
0.0     1.0     0.0     0.0
0.725   0.003   0.049   0.223
<
>pwms_22        8
0.428   0.513   0.003   0.055
0.839   0.038   0.002   0.121
0.427   0.092   0.104   0.378
0.094   0.097   0.637   0.171
0.78    0.012   0.023   0.185
0.907   0.007   0.064   0.022
0.0     0.121   0.0     0.879
0.063   0.352   0.088   0.498
<
>pwms_23        8
0.219   0.738   0.016   0.027
0.365   0.535   0.051   0.049
0.04    0.954   0.0     0.007
0.027   0.043   0.923   0.008
0.151   0.789   0.028   0.032
0.001   0.803   0.083   0.113
0.475   0.008   0.47    0.047
0.483   0.32    0.049   0.148
<
>pwms_24        8
0.455   0.15    0.065   0.331
0.379   0.162   0.247   0.212
0.124   0.034   0.065   0.777
0.001   0.205   0.0     0.794
0.038   0.052   0.059   0.851
0.215   0.02    0.603   0.162
0.961   0.022   0.014   0.003
0.971   0.0     0.027   0.002
<
```

```
>pwms_25	8
0.06	0.236	0.689	0.015
0.724	0.249	0.0	0.027
0.18	0.635	0.01	0.175
0.0	0.0	0.047	0.953
0.067	0.408	0.012	0.514
0.012	0.0	0.059	0.929
0.041	0.542	0.016	0.401
0.03	0.835	0.037	0.099
<
>pwms_26	8
0.135	0.017	0.211	0.637
0.196	0.127	0.311	0.367
0.022	0.06	0.902	0.016
0.006	0.167	0.68	0.147
0.017	0.951	0.011	0.021
0.008	0.009	0.948	0.035
0.004	0.343	0.647	0.006
0.003	0.161	0.825	0.01
<
>pwms_27	8
0.731	0.005	0.007	0.258
0.083	0.001	0.799	0.117
0.0	0.652	0.024	0.324
0.011	0.006	0.11	0.873
0.013	0.011	0.929	0.046
0.055	0.173	0.011	0.761
0.066	0.459	0.241	0.234
0.689	0.114	0.004	0.194
<
>pwms_28	8
0.064	0.436	0.0	0.499
0.075	0.007	0.003	0.916
0.054	0.419	0.524	0.003
0.051	0.543	0.047	0.359
0.092	0.0	0.0	0.908
0.019	0.458	0.521	0.002
0.157	0.842	0.0	0.002
0.055	0.02	0.0	0.926
<
>pwms_29	8
0.644	0.03	0.321	0.005
0.754	0.109	0.128	0.009
0.35	0.278	0.331	0.041
0.07	0.84	0.059	0.03
0.446	0.032	0.401	0.121
0.034	0.961	0.005	0.0
0.098	0.0	0.843	0.059
0.087	0.207	0.033	0.673
<
```

```
>pwms_30	8
0.976	0.0	0.024	0.0
0.848	0.012	0.066	0.073
0.115	0.63	0.255	0.0
0.448	0.296	0.25	0.005
0.001	0.208	0.776	0.015
0.002	0.553	0.276	0.169
0.106	0.644	0.236	0.015
0.88	0.001	0.001	0.118
<
>pwms_31	8
0.564	0.0	0.326	0.11
0.54	0.001	0.406	0.053
0.79	0.193	0.013	0.004
0.0	0.008	0.93	0.062
0.061	0.008	0.058	0.873
0.055	0.009	0.771	0.165
0.647	0.108	0.2	0.044
0.744	0.009	0.206	0.04
<
>pwms_32	8
0.002	0.0	0.761	0.237
0.026	0.021	0.239	0.714
0.147	0.178	0.617	0.058
0.709	0.182	0.0	0.109
0.002	0.932	0.01	0.056
0.931	0.058	0.001	0.011
0.13	0.002	0.865	0.003
0.153	0.432	0.097	0.319
<
>pwms_33	8
0.346	0.374	0.073	0.208
0.671	0.014	0.2	0.115
0.009	0.822	0.169	0.001
0.0	0.05	0.937	0.013
0.076	0.677	0.006	0.24
0.023	0.122	0.818	0.037
0.017	0.004	0.913	0.066
0.074	0.121	0.3	0.505
<
>pwms_34	8
0.766	0.063	0.0	0.171
0.094	0.152	0.228	0.526
0.01	0.052	0.912	0.026
0.911	0.023	0.014	0.053
0.355	0.33	0.312	0.003
0.191	0.023	0.086	0.701
0.104	0.125	0.012	0.759
0.195	0.067	0.003	0.735
<
```

```
>pwms_35        8
0.0     0.003   0.966   0.03
0.996   0.0     0.003   0.001
0.952   0.0     0.046   0.001
0.007   0.482   0.314   0.197
0.027   0.378   0.0     0.595
0.221   0.028   0.091   0.66
0.239   0.466   0.264   0.031
0.281   0.243   0.42    0.056
<
>pwms_36        8
0.055   0.628   0.317   0.0
0.0     0.0     0.038   0.961
0.053   0.141   0.539   0.267
0.632   0.001   0.367   0.0
0.01    0.836   0.002   0.152
0.228   0.772   0.0     0.001
0.0     0.378   0.005   0.617
0.042   0.0     0.252   0.706
<
>pwms_37        8
0.271   0.453   0.167   0.11
0.296   0.684   0.008   0.012
0.614   0.0     0.004   0.382
0.0     0.002   0.997   0.001
0.0     0.006   0.808   0.187
0.0     0.119   0.274   0.607
0.019   0.64    0.153   0.187
0.967   0.018   0.0     0.015
<
>pwms_38        8
0.128   0.667   0.04    0.165
0.107   0.892   0.0     0.002
0.82    0.0     0.001   0.179
0.0     0.005   0.992   0.003
0.016   0.011   0.966   0.007
0.222   0.332   0.211   0.234
0.0     0.737   0.162   0.101
0.71    0.0     0.279   0.01
<
>pwms_39        8
0.847   0.01    0.0     0.142
0.094   0.882   0.007   0.017
0.001   0.106   0.141   0.753
0.144   0.023   0.019   0.814
0.394   0.443   0.163   0.0
0.315   0.347   0.126   0.213
0.729   0.068   0.202   0.0
0.76    0.006   0.234   0.0
<
```

```
>pwms_40        8
0.117   0.0     0.871   0.013
0.002   0.29    0.021   0.687
0.021   0.411   0.264   0.304
0.068   0.12    0.036   0.776
0.0     0.049   0.898   0.053
0.0     0.201   0.002   0.797
0.5     0.041   0.26    0.199
0.717   0.002   0.031   0.251
<
>pwms_41        8
0.561   0.01    0.413   0.016
0.423   0.04    0.052   0.486
0.385   0.382   0.084   0.149
0.054   0.477   0.469   0.001
0.104   0.168   0.36    0.368
0.0     0.927   0.073   0.0
0.065   0.003   0.932   0.0
0.026   0.001   0.228   0.745
<
>pwms_42        8
0.071   0.302   0.002   0.625
0.177   0.051   0.056   0.717
0.022   0.937   0.0     0.041
0.51    0.371   0.048   0.071
0.0     0.769   0.002   0.229
0.005   0.001   0.971   0.023
0.13    0.733   0.04    0.098
0.029   0.443   0.417   0.11
<
>pwms_43        8
0.042   0.694   0.024   0.239
0.137   0.857   0.003   0.002
0.795   0.199   0.002   0.004
0.578   0.357   0.017   0.048
0.052   0.043   0.01    0.895
0.124   0.763   0.018   0.095
0.695   0.14    0.018   0.147
0.217   0.116   0.105   0.562
<
>pwms_44        8
0.003   0.005   0.001   0.992
0.013   0.01    0.0     0.976
0.19    0.448   0.0     0.363
0.545   0.02    0.341   0.094
0.739   0.045   0.208   0.008
0.692   0.082   0.12    0.106
0.221   0.122   0.034   0.623
0.153   0.07    0.111   0.666
<
```

>pwms_45 8
0.033 0.246 0.672 0.049
0.007 0.247 0.005 0.741
0.374 0.003 0.6 0.023
0.0 0.362 0.208 0.429
0.374 0.001 0.61 0.015
0.07 0.918 0.01 0.003
0.319 0.037 0.64 0.005
0.0 0.924 0.044 0.032
<
>pwms_46 8
0.189 0.473 0.334 0.003
0.632 0.017 0.321 0.03
0.649 0.106 0.006 0.239
0.01 0.001 0.986 0.003
0.0 0.0 0.001 0.999
0.0 0.073 0.425 0.502
0.009 0.824 0.166 0.001
0.375 0.053 0.236 0.336
<
>pwms_47 8
0.888 0.002 0.071 0.039
0.277 0.56 0.017 0.146
0.167 0.305 0.517 0.011
0.11 0.077 0.687 0.126
0.01 0.918 0.072 0.0
0.004 0.014 0.972 0.011
0.07 0.068 0.34 0.523
0.128 0.04 0.387 0.445
<
>pwms_48 8
0.95 0.016 0.033 0.002
0.707 0.182 0.004 0.107
0.831 0.085 0.051 0.032
0.195 0.446 0.354 0.005
0.059 0.169 0.771 0.001
0.003 0.966 0.031 0.001
0.157 0.084 0.465 0.294
0.049 0.054 0.75 0.147
<
>pwms_49 8
0.319 0.109 0.151 0.421
0.475 0.056 0.017 0.452
0.648 0.01 0.204 0.138
0.003 0.002 0.448 0.547
0.733 0.172 0.059 0.036
0.931 0.067 0.002 0.0
0.008 0.172 0.087 0.733
0.185 0.001 0.014 0.8
<

```
>pwms_50	8
0.017	0.54	0.051	0.392
0.605	0.382	0.008	0.004
0.138	0.014	0.122	0.726
0.0	0.371	0.618	0.011
0.015	0.101	0.0	0.884
0.101	0.092	0.694	0.114
0.535	0.045	0.013	0.408
0.0	0.076	0.082	0.842
<
>pwms_51	8
0.029	0.0	0.407	0.564
0.306	0.013	0.002	0.68
0.6	0.367	0.0	0.033
0.245	0.096	0.308	0.351
0.294	0.0	0.002	0.704
0.168	0.021	0.677	0.134
0.535	0.0	0.0	0.465
0.002	0.011	0.986	0.001
<
>pwms_52	8
0.137	0.221	0.158	0.483
0.378	0.023	0.013	0.585
0.004	0.029	0.967	0.0
0.003	0.145	0.001	0.851
0.001	0.0	0.0	0.999
0.104	0.01	0.459	0.428
0.385	0.078	0.055	0.482
0.08	0.538	0.015	0.367
<
>pwms_53	8
0.029	0.11	0.853	0.008
0.014	0.91	0.001	0.075
0.049	0.018	0.901	0.033
0.011	0.93	0.046	0.013
0.018	0.1	0.839	0.043
0.154	0.316	0.21	0.32
0.005	0.055	0.505	0.435
0.063	0.296	0.146	0.495
<
>pwms_54	8
0.183	0.01	0.576	0.232
0.0	0.0	0.906	0.094
0.002	0.028	0.589	0.381
0.035	0.022	0.653	0.29
0.122	0.275	0.104	0.499
0.0	0.783	0.171	0.046
0.995	0.001	0.004	0.0
0.001	0.0	0.999	0.0
<
```

```
>pwms_55	8
0.08	0.001	0.884	0.035
0.033	0.0	0.909	0.058
0.684	0.097	0.219	0.0
0.992	0.0	0.0	0.008
0.001	0.24	0.355	0.404
0.0	0.289	0.028	0.682
0.175	0.044	0.578	0.202
0.721	0.115	0.085	0.079
<
>pwms_56	8
0.0	0.918	0.08	0.002
0.972	0.002	0.003	0.023
0.206	0.057	0.62	0.117
0.001	0.695	0.087	0.217
0.141	0.143	0.085	0.63
0.014	0.617	0.342	0.027
0.108	0.04	0.067	0.785
0.0	0.0	0.997	0.002
<
>pwms_57	8
0.002	0.118	0.382	0.498
0.127	0.02	0.043	0.81
0.001	0.038	0.0	0.961
0.067	0.92	0.006	0.006
0.444	0.094	0.103	0.359
0.332	0.45	0.065	0.152
0.28	0.0	0.009	0.711
0.051	0.017	0.096	0.836
<
>pwms_58	8
0.78	0.002	0.051	0.167
0.581	0.032	0.059	0.328
0.025	0.015	0.191	0.769
0.07	0.171	0.088	0.672
0.219	0.227	0.547	0.007
0.581	0.014	0.126	0.279
0.998	0.0	0.0	0.002
0.785	0.01	0.191	0.014
<
>pwms_59	8
0.003	0.956	0.005	0.036
0.012	0.819	0.164	0.005
0.086	0.0	0.914	0.0
0.115	0.658	0.133	0.094
0.022	0.939	0.009	0.03
0.17	0.001	0.623	0.206
0.168	0.623	0.057	0.152
0.071	0.12	0.753	0.057
<
```

```
>pwms_60        8
0.0     0.012   0.14    0.848
0.038   0.776   0.138   0.047
0.867   0.052   0.001   0.08
0.034   0.708   0.182   0.076
0.814   0.14    0.04    0.006
0.283   0.079   0.298   0.34
0.146   0.761   0.028   0.065
0.065   0.934   0.001   0.0
<
>pwms_61        8
0.106   0.002   0.123   0.768
0.348   0.044   0.018   0.591
0.302   0.0     0.105   0.593
0.047   0.015   0.039   0.899
0.122   0.006   0.873   0.0
0.624   0.0     0.359   0.017
0.61    0.201   0.141   0.047
0.604   0.003   0.037   0.357
<
>pwms_62        8
0.031   0.192   0.468   0.309
0.007   0.065   0.122   0.806
0.05    0.144   0.744   0.061
0.877   0.05    0.005   0.068
0.175   0.658   0.018   0.149
0.894   0.005   0.1     0.0
0.655   0.04    0.305   0.0
0.29    0.7     0.011   0.0
<
>pwms_63        8
0.273   0.544   0.174   0.009
0.212   0.759   0.017   0.013
0.002   0.997   0.0     0.001
0.0     0.912   0.001   0.087
0.103   0.741   0.114   0.042
0.0     0.976   0.002   0.022
0.531   0.0     0.018   0.452
0.024   0.548   0.425   0.004
<
>pwms_64        8
0.0     0.304   0.0     0.696
0.101   0.443   0.295   0.161
0.801   0.128   0.063   0.008
0.821   0.173   0.003   0.004
0.692   0.033   0.038   0.237
0.0     0.867   0.133   0.0
0.002   0.106   0.279   0.612
0.2     0.766   0.023   0.011
<
```

```
>pwms_65	8
0.0	0.239	0.682	0.079
0.904	0.001	0.001	0.094
0.993	0.0	0.007	0.0
0.479	0.469	0.051	0.002
0.0	0.35	0.224	0.425
0.024	0.13	0.008	0.838
0.312	0.52	0.168	0.0
0.025	0.4	0.115	0.46
<
>pwms_66	8
0.104	0.0	0.033	0.863
0.015	0.876	0.099	0.01
0.013	0.947	0.041	0.0
0.387	0.347	0.0	0.266
0.092	0.005	0.842	0.061
0.122	0.041	0.819	0.017
0.136	0.0	0.847	0.017
0.0	0.305	0.676	0.019
<
>pwms_67	8
0.027	0.884	0.085	0.003
0.0	0.0	0.13	0.87
0.318	0.014	0.661	0.007
0.008	0.026	0.493	0.473
0.003	0.967	0.0	0.03
0.118	0.0	0.003	0.879
0.0	0.172	0.732	0.096
0.119	0.309	0.159	0.413
<
>pwms_68	8
0.0	0.879	0.121	0.0
0.323	0.001	0.084	0.592
0.072	0.0	0.003	0.925
0.033	0.099	0.022	0.847
0.182	0.569	0.052	0.197
0.479	0.02	0.459	0.042
0.444	0.086	0.013	0.457
0.533	0.36	0.004	0.103
<
>pwms_69	8
0.0	0.224	0.024	0.752
0.012	0.005	0.073	0.911
0.108	0.333	0.239	0.32
0.72	0.144	0.071	0.064
0.346	0.0	0.654	0.0
0.398	0.498	0.102	0.002
0.124	0.057	0.0	0.819
0.001	0.079	0.0	0.92
<
```

```
>pwms_70	8
0.89	0.015	0.021	0.074
0.218	0.617	0.163	0.002
0.706	0.293	0.0	0.001
0.046	0.948	0.0	0.006
0.037	0.313	0.004	0.647
0.153	0.004	0.154	0.69
0.025	0.005	0.907	0.063
0.067	0.386	0.238	0.309
<
>pwms_71	8
0.591	0.242	0.0	0.167
0.068	0.0	0.438	0.494
0.002	0.006	0.955	0.036
0.004	0.358	0.016	0.622
0.456	0.009	0.174	0.361
0.201	0.11	0.0	0.689
0.16	0.053	0.004	0.784
0.049	0.856	0.008	0.087
<
>pwms_72	8
0.69	0.003	0.288	0.019
0.003	0.836	0.148	0.013
0.976	0.0	0.012	0.012
0.392	0.0	0.598	0.01
0.28	0.58	0.108	0.033
0.147	0.0	0.075	0.779
0.179	0.052	0.769	0.0
0.07	0.09	0.738	0.102
<
>pwms_73	8
0.596	0.11	0.247	0.047
0.044	0.713	0.023	0.22
0.021	0.0	0.059	0.92
0.002	0.033	0.954	0.01
0.23	0.0	0.392	0.378
0.099	0.75	0.041	0.11
0.826	0.16	0.013	0.001
0.348	0.587	0.001	0.065
<
>pwms_74	8
0.263	0.027	0.0	0.71
0.169	0.476	0.155	0.2
0.24	0.004	0.001	0.755
0.0	0.011	0.055	0.933
0.015	0.238	0.499	0.249
0.374	0.014	0.544	0.069
0.826	0.067	0.103	0.004
0.925	0.0	0.002	0.073
<
```

```
>pwms_75	8
0.1	0.026	0.0	0.874
0.007	0.0	0.569	0.424
0.0	0.0	0.666	0.334
0.308	0.083	0.0	0.609
0.554	0.362	0.017	0.067
0.514	0.002	0.001	0.484
0.0	0.219	0.781	0.0
0.189	0.716	0.014	0.081
<
>pwms_76	8
0.008	0.928	0.023	0.041
0.813	0.0	0.147	0.039
0.0	0.391	0.034	0.575
0.151	0.002	0.845	0.003
0.144	0.156	0.003	0.697
0.0	0.071	0.779	0.15
0.0	0.13	0.668	0.203
0.278	0.425	0.201	0.097
<
>pwms_77	8
0.302	0.361	0.0	0.337
0.818	0.0	0.182	0.0
0.01	0.068	0.616	0.307
0.037	0.001	0.962	0.0
0.372	0.151	0.01	0.468
0.0	0.0	0.991	0.009
0.006	0.442	0.003	0.549
0.471	0.007	0.317	0.205
<
>pwms_78	8
0.285	0.614	0.101	0.001
0.615	0.26	0.038	0.088
0.305	0.007	0.201	0.487
0.019	0.508	0.167	0.306
0.0	0.084	0.289	0.627
0.868	0.082	0.048	0.002
0.947	0.007	0.003	0.043
0.935	0.064	0.0	0.001
<
>pwms_79	8
0.057	0.031	0.765	0.147
0.0	0.127	0.558	0.316
0.219	0.064	0.684	0.033
0.001	0.0	0.003	0.996
0.327	0.018	0.323	0.332
0.0	0.0	0.848	0.152
0.069	0.623	0.214	0.094
0.003	0.965	0.0	0.032
<
```

```
>pwms_80	8
0.503	0.083	0.411	0.003
0.017	0.941	0.001	0.042
0.817	0.037	0.033	0.113
0.0	0.022	0.606	0.372
0.684	0.177	0.017	0.122
0.0	0.486	0.409	0.105
0.084	0.857	0.01	0.048
0.0	0.158	0.0	0.842
<
>pwms_81	8
0.033	0.551	0.333	0.084
0.0	0.698	0.01	0.292
0.001	0.988	0.01	0.001
0.0	0.878	0.02	0.102
0.142	0.004	0.067	0.787
0.003	0.015	0.977	0.005
0.099	0.037	0.862	0.001
0.463	0.031	0.394	0.112
<
>pwms_82	8
0.564	0.002	0.163	0.271
0.579	0.033	0.094	0.294
0.023	0.031	0.933	0.012
0.002	0.033	0.03	0.934
0.018	0.19	0.485	0.307
0.0	0.994	0.005	0.0
0.424	0.022	0.007	0.547
0.119	0.302	0.07	0.509
<
>pwms_83	8
0.522	0.005	0.038	0.435
0.232	0.084	0.684	0.0
0.726	0.088	0.024	0.161
0.063	0.847	0.0	0.089
0.935	0.055	0.002	0.007
0.53	0.009	0.424	0.036
0.245	0.671	0.004	0.08
0.31	0.005	0.0	0.685
<
>pwms_84	8
0.358	0.259	0.211	0.172
0.112	0.816	0.021	0.051
0.135	0.158	0.661	0.045
0.07	0.801	0.014	0.114
0.095	0.001	0.876	0.028
0.41	0.057	0.509	0.024
0.013	0.965	0.021	0.002
0.149	0.023	0.826	0.002
<
```

>pwms_85	8
0.49	0.069	0.003	0.438
0.25	0.0	0.674	0.076
0.193	0.394	0.002	0.412
0.004	0.127	0.001	0.868
0.018	0.001	0.946	0.035
0.088	0.034	0.0	0.879
0.002	0.725	0.01	0.263
0.442	0.342	0.065	0.151
<
>pwms_86	8
0.217	0.023	0.661	0.1
0.001	0.954	0.001	0.045
0.021	0.439	0.539	0.0
0.083	0.015	0.694	0.208
0.225	0.001	0.756	0.017
0.902	0.04	0.0	0.057
0.832	0.01	0.155	0.003
0.246	0.061	0.386	0.307
<
>pwms_87	8
0.216	0.342	0.031	0.411
0.001	0.396	0.016	0.587
0.338	0.024	0.007	0.632
0.173	0.817	0.009	0.001
0.646	0.285	0.058	0.01
0.052	0.806	0.141	0.0
0.0	0.0	0.824	0.176
0.171	0.806	0.012	0.011
<
>pwms_88	8
0.726	0.05	0.096	0.128
0.073	0.24	0.028	0.659
0.001	0.036	0.001	0.962
0.016	0.001	0.027	0.956
0.074	0.611	0.087	0.229
0.593	0.066	0.071	0.27
0.77	0.2	0.021	0.008
0.414	0.003	0.111	0.472
<
>pwms_89	8
0.28	0.0	0.718	0.002
0.023	0.707	0.27	0.0
0.984	0.001	0.002	0.013
0.011	0.956	0.0	0.033
0.272	0.003	0.0	0.725
0.1	0.425	0.435	0.039
0.729	0.126	0.134	0.011
0.268	0.164	0.227	0.341
<

```
>pwms_90	8
0.752	0.091	0.047	0.109
0.016	0.599	0.0	0.385
0.038	0.112	0.848	0.001
0.132	0.337	0.479	0.052
0.001	0.925	0.074	0.001
0.002	0.006	0.981	0.011
0.118	0.073	0.44	0.369
0.131	0.35	0.175	0.345
<
>pwms_91	8
0.141	0.1	0.202	0.558
0.056	0.029	0.652	0.263
0.41	0.019	0.006	0.565
0.41	0.278	0.001	0.311
0.975	0.0	0.001	0.024
0.682	0.157	0.161	0.0
0.016	0.964	0.002	0.017
0.758	0.113	0.021	0.109
<
>pwms_92	8
0.001	0.672	0.098	0.228
0.009	0.001	0.981	0.01
0.0	0.438	0.379	0.184
0.007	0.304	0.494	0.195
0.014	0.066	0.046	0.873
0.171	0.142	0.001	0.687
0.0	0.001	0.111	0.888
0.293	0.324	0.003	0.38
<
>pwms_93	8
0.133	0.762	0.017	0.088
0.0	1.0	0.0	0.0
0.556	0.297	0.143	0.004
0.702	0.011	0.286	0.001
0.086	0.0	0.914	0.0
0.272	0.026	0.676	0.027
0.001	0.851	0.106	0.042
0.54	0.001	0.004	0.455
<
>pwms_94	8
0.019	0.925	0.056	0.0
0.03	0.188	0.703	0.08
0.135	0.137	0.194	0.534
0.404	0.035	0.042	0.52
0.42	0.52	0.035	0.024
0.164	0.168	0.378	0.29
0.02	0.97	0.009	0.0
0.055	0.13	0.811	0.004
<
```

```
>pwms_95	8
0.093	0.817	0.015	0.074
0.002	0.003	0.982	0.013
0.176	0.105	0.035	0.683
0.0	0.143	0.853	0.004
0.009	0.115	0.765	0.111
0.362	0.058	0.046	0.534
0.077	0.521	0.176	0.227
0.098	0.746	0.154	0.003
<
>pwms_96	8
0.052	0.561	0.001	0.386
0.012	0.083	0.263	0.642
0.07	0.226	0.234	0.47
0.114	0.0	0.273	0.613
0.0	0.0	0.002	0.998
0.231	0.049	0.5	0.219
0.0	0.001	0.944	0.056
0.022	0.942	0.035	0.0
<
>pwms_97	8
0.008	0.984	0.0	0.008
0.001	0.008	0.968	0.023
0.445	0.277	0.266	0.012
0.142	0.227	0.515	0.115
0.697	0.124	0.11	0.069
0.677	0.291	0.016	0.016
0.306	0.139	0.389	0.166
0.64	0.0	0.0	0.36
<
>pwms_98	8
0.674	0.193	0.132	0.0
0.358	0.032	0.213	0.398
0.697	0.257	0.046	0.0
0.429	0.561	0.009	0.001
0.593	0.0	0.0	0.407
0.0	0.001	0.0	0.999
0.0	0.67	0.113	0.217
0.534	0.006	0.006	0.454
<
>pwms_99	8
0.617	0.011	0.356	0.015
0.066	0.113	0.082	0.739
0.021	0.002	0.405	0.572
0.168	0.817	0.0	0.015
0.201	0.053	0.021	0.726
0.101	0.086	0.022	0.791
0.25	0.242	0.014	0.493
0.004	0.082	0.009	0.904
<
```

```
>pwms_100	8
0.42	0.014	0.13	0.436
0.227	0.255	0.517	0.001
0.831	0.001	0.013	0.154
0.996	0.0	0.002	0.003
0.696	0.0	0.006	0.298
0.65	0.241	0.068	0.041
0.034	0.197	0.0	0.769
0.637	0.184	0.041	0.138
<
>pwms_101	8
0.27	0.136	0.439	0.155
0.0	0.993	0.0	0.007
0.0	0.0	0.043	0.957
0.006	0.021	0.969	0.003
0.422	0.013	0.399	0.166
0.098	0.596	0.067	0.239
0.332	0.643	0.005	0.02
0.176	0.823	0.0	0.0
<
>pwms_102	8
0.056	0.001	0.703	0.24
0.0	0.012	0.876	0.112
0.159	0.059	0.699	0.083
0.0	0.721	0.058	0.22
0.348	0.027	0.116	0.509
0.0	0.944	0.021	0.035
0.779	0.002	0.183	0.036
0.0	0.0	1.0	0.0
<
>pwms_103	8
0.004	0.709	0.111	0.176
0.023	0.095	0.396	0.486
0.012	0.003	0.135	0.849
0.173	0.431	0.151	0.245
0.041	0.887	0.041	0.031
0.005	0.683	0.247	0.065
0.003	0.04	0.953	0.004
0.079	0.91	0.0	0.011
<
>pwms_104	8
0.006	0.977	0.003	0.013
0.803	0.052	0.144	0.001
0.001	0.76	0.006	0.233
0.044	0.001	0.954	0.001
0.271	0.604	0.015	0.111
0.024	0.424	0.377	0.175
0.044	0.263	0.218	0.475
0.232	0.189	0.006	0.573
<
```

```
>pwms_105	8
0.997	0.002	0.001	0.0
0.009	0.443	0.0	0.548
0.0	0.542	0.211	0.247
0.135	0.002	0.161	0.701
0.011	0.479	0.004	0.507
0.012	0.003	0.717	0.268
0.506	0.301	0.193	0.0
0.546	0.028	0.426	0.0
<
>pwms_106	8
0.187	0.0	0.812	0.001
0.036	0.0	0.964	0.0
0.0	0.145	0.849	0.007
0.024	0.396	0.58	0.0
0.0	0.008	0.0	0.992
0.005	0.271	0.724	0.0
0.195	0.025	0.459	0.321
0.0	0.004	0.745	0.25
<
>pwms_107	8
0.142	0.0	0.131	0.727
0.221	0.027	0.663	0.089
0.069	0.013	0.062	0.857
0.185	0.534	0.014	0.267
0.816	0.177	0.0	0.007
0.702	0.166	0.047	0.085
0.005	0.943	0.051	0.0
0.491	0.348	0.005	0.156
<
>pwms_108	8
0.579	0.0	0.364	0.057
0.17	0.104	0.703	0.022
0.0	0.27	0.521	0.21
0.0	0.979	0.0	0.02
0.053	0.072	0.642	0.233
0.12	0.0	0.673	0.207
0.584	0.0	0.007	0.409
0.878	0.013	0.109	0.001
<
>pwms_109	8
0.359	0.001	0.284	0.356
0.462	0.0	0.267	0.271
0.002	0.094	0.436	0.468
0.749	0.169	0.017	0.065
0.897	0.001	0.101	0.001
0.091	0.241	0.001	0.667
0.08	0.0	0.28	0.64
0.006	0.0	0.002	0.992
<
```

```
>pwms_110	8
0.089	0.001	0.45	0.46
0.0	0.004	0.004	0.991
0.082	0.004	0.301	0.613
0.078	0.049	0.562	0.311
0.001	0.504	0.346	0.148
0.181	0.721	0.01	0.088
0.001	0.228	0.719	0.051
0.008	0.019	0.973	0.0
<
>pwms_111	8
0.802	0.006	0.003	0.189
0.257	0.143	0.007	0.593
0.019	0.074	0.068	0.839
0.008	0.784	0.206	0.002
0.248	0.689	0.05	0.012
0.0	0.019	0.006	0.975
0.176	0.034	0.383	0.407
0.543	0.19	0.054	0.212
<
>pwms_112	8
0.001	0.406	0.108	0.485
0.001	0.655	0.001	0.343
0.395	0.208	0.0	0.397
0.671	0.28	0.005	0.044
0.537	0.061	0.01	0.392
0.695	0.211	0.014	0.08
0.141	0.767	0.028	0.064
0.001	0.103	0.883	0.014
<
>pwms_113	8
0.048	0.0	0.457	0.494
0.003	0.961	0.036	0.0
0.741	0.127	0.0	0.132
0.202	0.001	0.77	0.027
0.208	0.09	0.694	0.008
0.018	0.039	0.373	0.57
0.001	0.848	0.13	0.02
0.093	0.013	0.033	0.861
<
>pwms_114	8
0.002	0.767	0.198	0.034
0.252	0.013	0.733	0.002
0.0	0.851	0.027	0.122
0.001	0.994	0.001	0.004
0.002	0.036	0.924	0.037
0.01	0.773	0.195	0.022
0.019	0.461	0.458	0.062
0.272	0.254	0.15	0.325
<
```

```
>pwms_115	8
0.538	0.0	0.04	0.422
0.203	0.015	0.775	0.007
0.065	0.011	0.866	0.059
0.048	0.033	0.844	0.075
0.033	0.053	0.585	0.329
0.094	0.725	0.154	0.027
0.061	0.935	0.004	0.001
0.0	0.194	0.0	0.806
<
>pwms_116	8
0.154	0.0	0.771	0.075
0.077	0.069	0.002	0.852
0.282	0.096	0.012	0.611
0.0	0.053	0.022	0.925
0.0	0.042	0.555	0.403
0.111	0.46	0.29	0.139
0.046	0.064	0.26	0.63
0.0	0.075	0.0	0.924
<
>pwms_117	8
0.009	0.051	0.533	0.407
0.031	0.815	0.013	0.141
0.0	0.473	0.504	0.023
0.063	0.204	0.522	0.212
0.212	0.019	0.649	0.121
0.156	0.625	0.007	0.212
0.0	0.997	0.0	0.003
0.003	0.033	0.957	0.007
<
>pwms_118	8
0.0	0.256	0.437	0.307
0.025	0.012	0.0	0.962
0.323	0.009	0.664	0.004
0.618	0.28	0.101	0.0
0.984	0.0	0.001	0.015
0.002	0.295	0.429	0.274
0.0	0.059	0.1	0.841
0.09	0.521	0.033	0.356
<
>pwms_119	8
0.168	0.173	0.646	0.014
0.233	0.09	0.607	0.069
0.115	0.649	0.048	0.188
0.036	0.729	0.235	0.0
0.008	0.051	0.941	0.0
0.096	0.0	0.89	0.014
0.844	0.15	0.006	0.001
0.788	0.122	0.088	0.002
<
```

```
>pwms_120	8
0.0	0.579	0.247	0.173
0.033	0.0	0.154	0.813
0.034	0.007	0.004	0.956
0.048	0.929	0.0	0.023
0.038	0.421	0.068	0.473
0.483	0.236	0.281	0.0
0.84	0.0	0.111	0.049
0.072	0.428	0.065	0.435
<
>pwms_121	8
0.988	0.005	0.001	0.006
0.015	0.539	0.001	0.445
0.0	0.358	0.012	0.629
0.04	0.007	0.006	0.946
0.033	0.0	0.496	0.471
0.105	0.102	0.143	0.65
0.223	0.083	0.484	0.21
0.011	0.519	0.013	0.457
<
>pwms_122	8
0.14	0.505	0.027	0.328
0.864	0.108	0.025	0.002
0.667	0.089	0.103	0.141
0.016	0.388	0.486	0.11
0.045	0.025	0.095	0.836
0.001	0.25	0.614	0.134
0.01	0.0	0.063	0.926
0.009	0.008	0.952	0.031
<
>pwms_123	8
0.809	0.0	0.086	0.104
0.257	0.01	0.0	0.732
0.057	0.034	0.907	0.002
0.714	0.153	0.0	0.132
0.747	0.001	0.172	0.081
0.122	0.446	0.005	0.427
0.0	0.039	0.335	0.626
0.239	0.029	0.481	0.251
<
>pwms_124	8
0.852	0.009	0.01	0.128
0.255	0.225	0.001	0.519
0.174	0.001	0.145	0.681
0.078	0.864	0.014	0.044
0.889	0.106	0.002	0.002
0.118	0.582	0.04	0.259
0.642	0.045	0.253	0.06
0.232	0.123	0.029	0.616
<
```

```
>pwms_125       8
0.375   0.525   0.005   0.095
0.082   0.905   0.0     0.013
0.667   0.221   0.003   0.109
0.175   0.0     0.825   0.0
0.044   0.668   0.143   0.145
0.929   0.001   0.067   0.004
0.009   0.0     0.991   0.0
0.0     0.278   0.665   0.056
<
>pwms_126       8
0.284   0.018   0.088   0.611
0.107   0.118   0.747   0.028
0.401   0.247   0.264   0.088
0.798   0.0     0.048   0.154
0.528   0.247   0.011   0.213
0.942   0.002   0.001   0.055
0.017   0.973   0.003   0.008
0.839   0.03    0.001   0.13
<
>pwms_127       8
0.145   0.631   0.148   0.076
0.335   0.123   0.336   0.206
0.125   0.842   0.005   0.027
0.0     0.011   0.944   0.045
0.028   0.378   0.02    0.574
0.0     0.007   0.989   0.004
0.003   0.192   0.709   0.097
0.211   0.103   0.0     0.685
<
```
