## Supplementary File for "SEAMoD: A fully interpretable neural network for cis-regulatory analysis of differentially expressed genes"

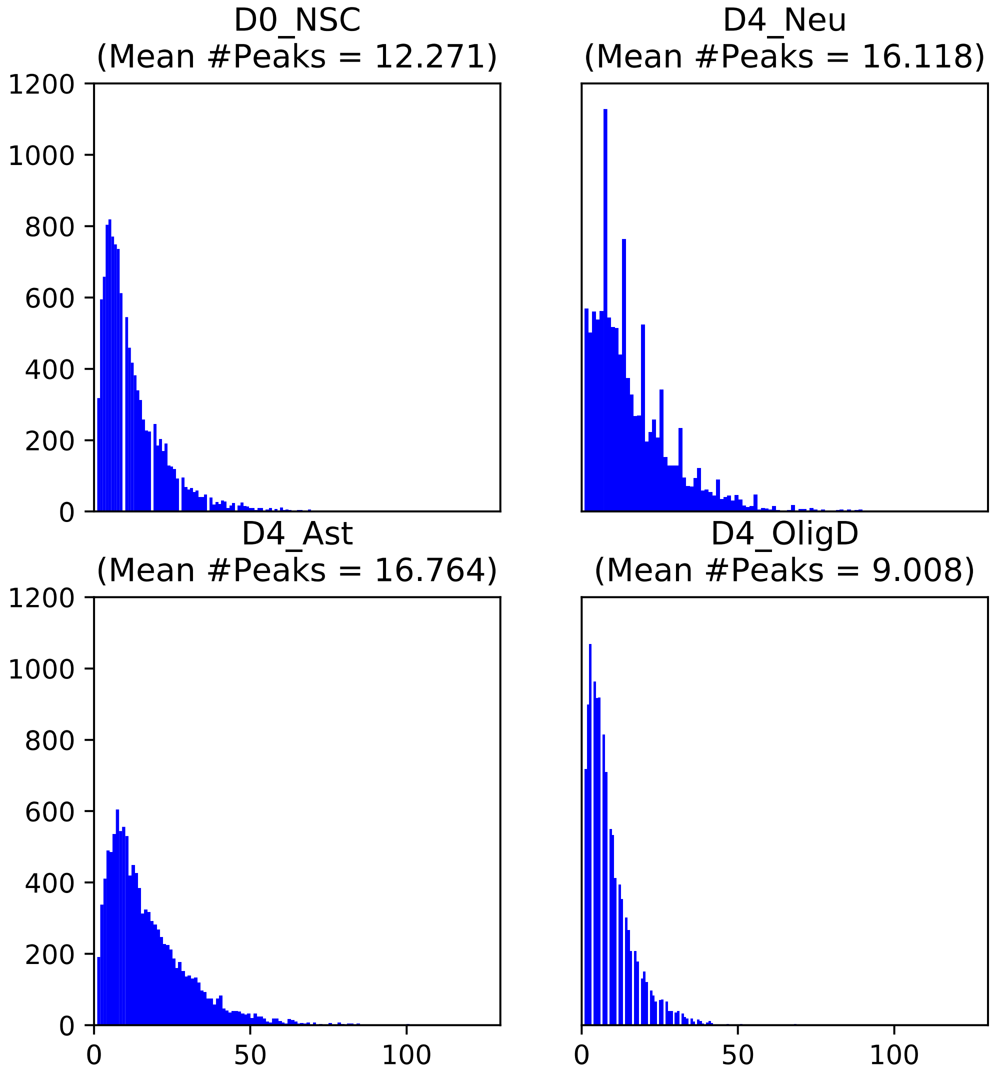


**Supplementary Figure 1: Number of peaks H3K27Ac peak in a gene's regulatory region.** Here each histogram represents a cell type with X axis indicating the number of peaks inside a gene's regulatory region. The regulatory region of a gene was defined as the 200kb window centered around the gene's transcription start site (TSS). Here, the number above a histogram indicates the average number of peaks across all genes in the corresponding cell type. Most of the genes had more than 10 peaks in their regulatory regions across all 4 cell types. Each one of these could have a regulatory impact on the gene.

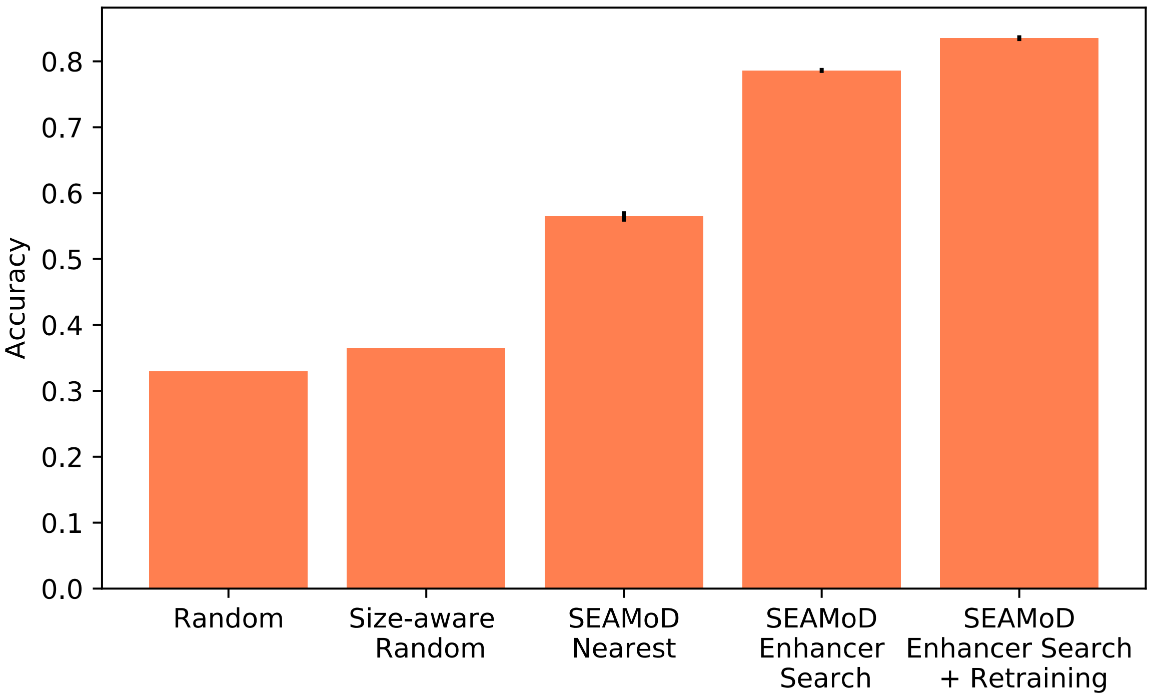


**Supplementary Figure 2: Model Performance on the seen dataset.** The bar plots show the model performance on the seen dataset which comprises of the genes that were used for training and validation. X axis represents the models (including the two random baselines – "Random" and "Size-aware" random) and Y axis represents the classification accuracy on the seen gene set (10,460 genes), averaged over ten folds, along with error bars (standard errors)

| D4_Neu | D4_Ast | D4_OligD | Number of Genes |
|---|---|---|---|
| -1 | -1 | -1 | 3430 |
| -1 | -1 | 0 | 467 |
| -1 | -1 | 1 | 62 |
| -1 | 0 | -1 | 189 |
| -1 | 0 | 0 | 382 |
| -1 | 0 | 1 | 122 |
| -1 | 1 | -1 | 18 |
| -1 | 1 | 0 | 34 |
| -1 | 1 | 1 | 92 |
| 0 | -1 | -1 | 431 |
| 0 | -1 | 0 | 193 |
| 0 | -1 | 1 | 31 |
| 0 | 0 | -1 | 266 |
| 0 | 0 | 0 | 0 |
| 0 | 0 | 1 | 340 |
| 0 | 1 | -1 | 26 |
| 0 | 1 | 0 | 331 |
| 0 | 1 | 1 | 552 |
| 1 | -1 | -1 | 53 |
| 1 | -1 | 0 | 19 |
| 1 | -1 | 1 | 25 |
| 1 | 0 | -1 | 89 |
| 1 | 0 | 0 | 440 |
| 1 | 0 | 1 | 218 |
| 1 | 1 | -1 | 51 |
| 1 | 1 | 0 | 499 |
| 1 | 1 | 1 | 3262 |

**Supplementary Table 1: Distribution of DEGs across the 3 target cell types.** Each gene can be divided into one of 27 classes (identified by the first 3 numbers in a row) represent by the 27 rows in this table. The numbers in the first three columns represent the DEG (Differentially Expressed Genes) class of the genes in the corresponding cell type (indicated by the column header) with -1, 0, and 1 being down-regulated, non-DE, and up-regulated respectively. The number in the 4$^{th}$ column gives the total number of genes in the corresponding DEG class. Most of the genes show same directionality in the gene expression changes across the three cell types as indicated by the first and the last rows.

| Peak Set | D0_NSC | D4_Neu | D4_Ast | D4_OligD |
|---|---|---|---|---|
| NN | 58 | 57 | 78 | 68 |
| BE | 136 | 128 | 139 | 135 |
| Total | 308 | 318 | 317 | 301 |

**Supplementary Table 2: Best enhancers show greater overlap with validated enhancers than the nearest neighbors do.** We tested the overlap between experimentally validated (EV) set of regulatory enhancers and the two sets of enhancers used in our study – nearest neighbors (NN) and the best enhancers (BE) identified by the model. The last row of the table represents the total number of genes for which there was an overlap between the EV enhancers of a gene and the nearest 4 H3K27Ac peaks in its regulatory region. The rows NN and BE represent the number of genes for which the nearest neighbor and the best enhancer overlapped with a gene's EV enhancer respectively. There is almost two-fold increase when we choose the best enhancer instead of the nearest neighbor across all 4 cell types. An H3K27Ac peak (BE or NN) and an EV enhancer is considered to overlap if they have at least 100 overlapping basepairs.

**Supplementary S1: Set of PWMs generated from the convolutional filters learnt by the model.** The file lists all 128 PWMs obtained from the convolutional filters by the process described in the methods. Note that the numbers have been rounded up to 3 significant digits for better readability and hence they might not add up to 1 at a few positions.
